# MERFISH-based pooled genetic screening reveals cell non-autonomous phenotypes in intact tissue

**DOI:** 10.64898/2026.10.06.756924

**Authors:** Kang Jin, Reuben A. Saunders, Jiahao Zhang, Jiaqi Lu, Aaron R. Halpern, Jaspreet Sandhu, Xingjie Pan, Cosmos Yuqi Wang, Alexander M. Li, William E. Allen, Jonathan S. Weissman, Xiaowei Zhuang

## Abstract

Genetic perturbations can reshape both the state of a targeted cell and the behavior of neighboring cells. Measuring these effects systematically in intact tissues requires pooled screening methods that jointly identify perturbations, molecular phenotypes, and spatial context of individual cells. Here, we report an imaging-based pooled screening method that detects perturbation identities and endogenous RNAs without enzymatic amplification. Relative to our previous enzymatic-amplification-based workflow, amplification-free MERFISH detection with direct hybridization increased recovery of perturbation barcodes by ∼10-fold and endogenous transcripts by 2.5-fold, improving perturbation assignment as well as molecular profiling of both perturbed and neighboring cells. Applying this method to a genetic mosaic mouse liver screen targeting ∼200 genes in hepatocytes, we profiled 1.5 million cells in intact tissues and mapped both cell-autonomous responses of perturbed hepatocytes and cell non-autonomous changes in neighboring immune and vascular populations. These included localized accumulation of pDC-like cells near hepatocytes with impaired V-ATPase function and angiogenic responses in vascular cells near *Vhl*-perturbed hepatocytes with hypoxia activation. Our method provides a simplified, scalable approach to linking genetic perturbations with cell state, multicellular response, and tissue organization.

## Introduction

Cells respond to genetic perturbations within structured tissues, where anatomical position, cellular morphology, and communication with neighboring cells shape the resulting phenotype. Measuring these dimensions at the scale of pooled genetic screening would enable systematic analysis of both cell-autonomous gene function and how perturbed cells reshape their local environments. Sequencing-based approaches such as Perturb-seq^1–4^ provide rich molecular readouts of genetic perturbations^5–7^, yet require dissociation of cells from tissues and extraction of molecules from cells, losing information about cellular morphology, anatomical position, and neighboring cells.

Imaging-based pooled screens connect perturbations to spatially resolved RNA, protein, and morphological phenotypes while preserving tissue context^8–19^. Advances in imaging-based spatial omics have enabled measurements of thousands of RNA species and dozens of proteins while retaining subcellular localization of molecules and spatial organization of cells in tissue^20–23^. A central challenge, however, is identifying the genetic perturbation carried by each imaged cell.

The short sequences used to identify perturbations, including sgRNAs and associated barcodes, provide limited space for probe binding. Previous imaging-based pooled screens have therefore used signal or target amplification. An early pooled CRISPR screen from our laboratory used long RNA barcodes (450 nucleotides (nt)) and branched-DNA signal amplification^24, 25^ to identify sgRNAs and screen for regulators of subcellular RNA localization in cultured cells^10^. Although effective for barcode detection, these long barcode sequences complicated the synthesis and pooled cloning of large perturbation libraries.

More recent methods use shorter sequences to facilitate pooled library construction, together with enzymatic amplification to help perturbation detection. Perturb-FISH^18^ and PerturbView^17^ use *in vitro* transcription^26^ to amplify perturbation-identifying RNAs, whereas CRISPRmap^15^, Perturb-Multimodal (Perturb-Multi)^16^, and RAEFISH^19^ use rolling-circle amplification (RCA)^27–29^ to amplify guide-associated barcodes or variable sequences in sgRNAs. In Perturb-Multi, we used 185-nt barcodes to support pooled library construction and amplified the barcode sequences by RCA before detection with multiplexed error-robust fluorescence *in situ* hybridization (MERFISH)^30^. Because RCA degrades encoding probes used in conventional MERFISH, endogenous mRNAs were also detected using RCA-MERFISH to maintain protocol compatibility. These enzymatic reactions add processing steps and can reduce the detection efficiency of barcode and/or endogenous RNAs, potentially limiting both perturbation assignment and the sensitivity for measuring responses in perturbed and neighboring cells.

We sought to overcome these limitations and asked whether the many copies of barcode RNA expressed in each infected cell would permit reliable perturbation assignment by direct hybridization without amplification, despite the limited number of probe-binding sites per barcode. Notably, we demonstrated robust MERFISH detection of barcodes as short as 90 nt without enzymatic amplification. In liver tissue, this enzymatic-amplification-free approach increased the detection efficiency of perturbation barcodes and endogenous transcripts per cell by ∼10-fold and 2.5-fold, respectively, relative to our previous RCA-MERFISH based workflow. Applying this amplification-free Perturb-MERFISH protocol (Fig. 1a) to a pooled CRISPR screen in mouse liver, we not only observed expected cell-autonomous responses, but also uncovered many cell non-autonomous effects of perturbed hepatocytes on the abundance and transcriptional states of neighboring immune and stromal populations. In particular, our cell non-autonomous analyses identified localized accumulation of plasmacytoid dendritic cell (pDC)–like cells near hepatocytes with impaired V-ATPase function and angiogenic responses in liver sinusoidal endothelial cells (LSECs) and hepatic stellate cells (HSCs) near *Vhl*-perturbed hepatocytes that activate a hypoxia program. This amplification-free Perturb-MERFISH approach thus enables systematic analysis of how perturbed cells reshape their local tissue environments.

**Fig. 1.**
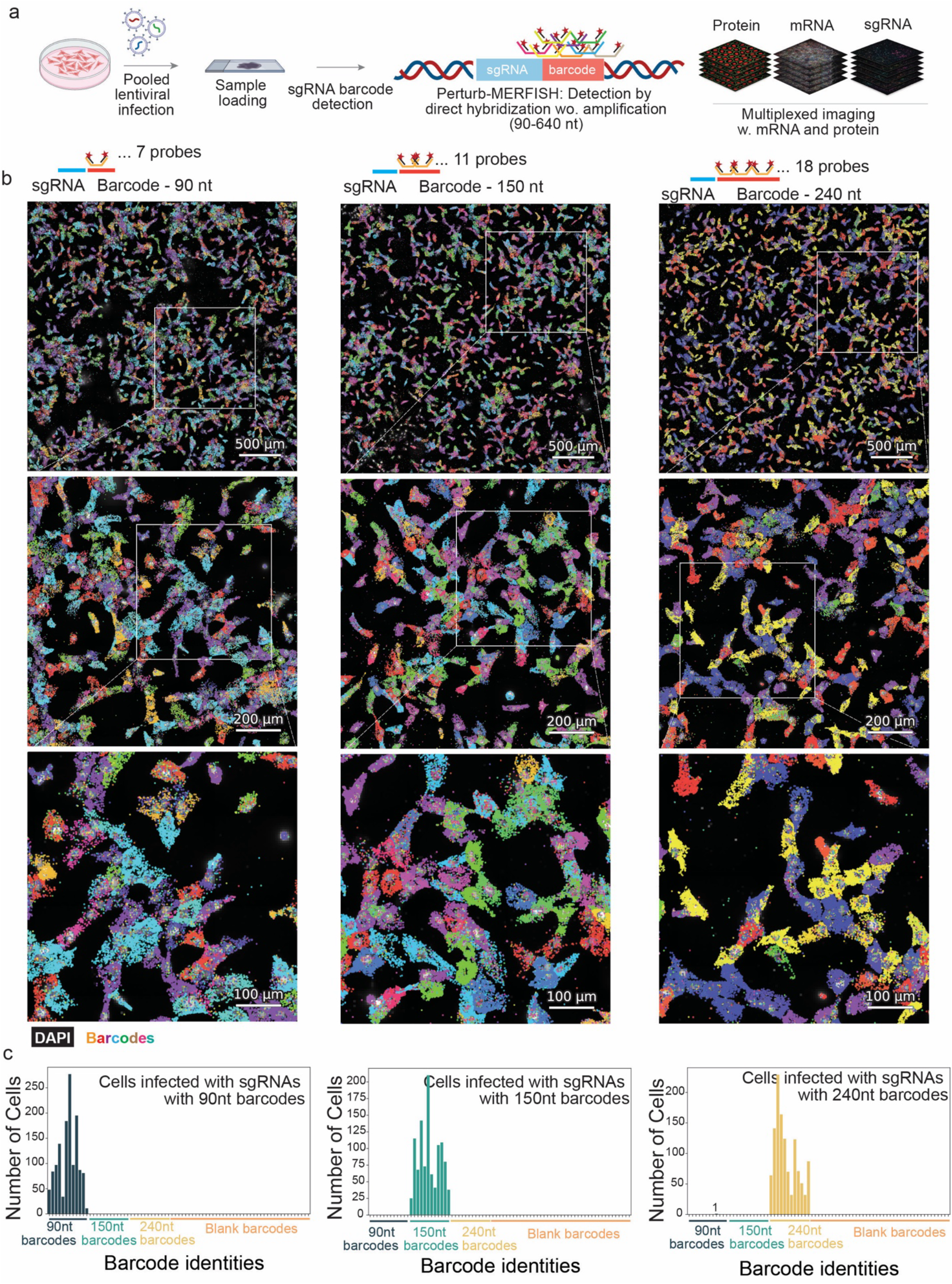
MERFISH-based *in situ* detection of genetic perturbation by amplification-free imaging of compact barcodes associated with sgRNA detection. **a,** Schematic of MERFISH-based pooled genetic screen by amplification-free, *in situ* sgRNA detection using compact barcodes, integrated with imaging of endogenous mRNAs and proteins. Some elements in this panel were created with BioRender. Jin, K. (2026) https://BioRender.com/yww91zb. **b,** Detection of sgRNAs using 90-, 150-, and 240-nt long barcodes in cultured RPE1 cells. Images show detected sgRNA-associated barcodes in different colors, overlaid with DAPI-stained nuclei. For each barcode length, a global field of view (FOV) is shown at the top, followed by two successive magnifications of the indicated regions in the middle and bottom rows. Schematics above each column show sgRNA in blue, associated barcodes in red, MERFISH encoding probes in yellow, and readout probes shown in black with stars indicating dyes. **c,** Assessment of barcode-detection specificity in RPE1 cells separately infected with sgRNA–barcode pools containing 90-, 150-, or 240-nt barcodes, but detected using a common MERFISH probe library targeting barcodes from all three pools. Assignment of cells to barcodes belonging to the two pools not added to the sample provides a direct measure of misidentification. Plots show the distribution of sgRNA–barcode identities assigned to segmented cells (Materials and Methods). For each infection condition, nearly all assigned cells were mapped to barcodes from the corresponding pool, with negligible assignments to barcodes from the other pools or to blank barcodes, demonstrating high specificity of sgRNA barcode detection.

## Results

### Amplification-free MERFISH detection of compact sgRNA barcodes *in vitro*

To test whether direct hybridization without target amplification could support sensitive and robust sgRNA-barcode detection, we established a pilot system in RPE1 cells^6^ and tested barcode designs ranging from 90 to 640 nt using a vector derived from a modified CROP-seq construct^31, 32^. Details of the vector construction and barcode design are described in Materials and Methods, Fig. 1a, and Supplementary Fig. 1a.

We transduced RPE1 cells at low multiplicity of infection (MOI) and selected infected cells with puromycin before performing MERFISH. We first evaluated a library of 124 sgRNA– barcode pairs with 640-nt barcodes, covering 56 target genes and six non-targeting controls. Direct hybridization to the 640-nt barcodes without target amplification produced bright, discrete puncta with low background across large imaging areas (Supplementary Fig. 1b). In contrast to endogenous transcripts, which were broadly distributed across cells, each perturbation barcode identity was concentrated within a small subset of cells, as expected from infection of individual cells predominantly by a single lentiviral construct at low MOI^33, 34^ (Supplementary Fig. 1c). Among cells classified as sgRNA singlets (cells receiving a single perturbation; see Materials and Methods section for classification criteria), the dominant barcode identity accounted for 87% of detected barcode molecules per cell, substantially higher than in ambiguous cells that could not be confidently classified as singlets or multiplets (cells receiving a combination of distinct perturbations; Supplementary Fig. 1d). For perturbed cells, decoding yielded a median of 193 detected barcode molecules corresponding to the assigned sgRNA per cell (Supplementary Fig. 1e), providing abundant signal for robust sgRNA assignment. These results established that expressed perturbation barcodes could be identified reliably by direct hybridization without amplification.

### Compact sgRNA barcodes support robust amplification-free detection *in vitro*

To facilitate pooled library construction and future *in vivo* applications, we next tested shorter barcodes. We tested barcodes of 90, 150, and 240 nt in length, which accommodated only 7, 11, and 18 MERFISH encoding probes, respectively (Fig. 1b). All three designs generated readily detectable, cell-restricted signals and supported unambiguous decoding across diverse fields of view (Fig. 1b). Testing the detection crosstalk by infecting the samples with a lentiviral library carrying one of the three barcode pools, but detecting the barcodes with a MERFISH encoding library targeting all pools simultaneously (Fig. 1c), we estimated a cross-library misidentification rate of 0.03% (Materials and Methods), demonstrating that direct hybridization without target amplification can identify perturbations with high specificity for barcodes as short as 90 nt.

### Amplification-free barcode detection allows pooled perturbation assignment *in vivo*

We next asked whether amplification-free direct sgRNA-barcode detection could identify pooled genetic perturbations in intact tissue. In a pilot experiment using a pooled lentiviral library containing 260-nt barcodes, direct hybridization without target amplification produced strong, cell-restricted signals with low background across large liver sections, enabling clear perturbation assignment within densely organized tissue (Supplementary Fig. 2a).

We then applied the workflow to perform pooled genetic screens in the mouse liver (Fig. 2a), using a pooled lentiviral CRISPR sgRNA perturbation library generated in our earlier study^16^ following the CROP-seq design^31^, perturbing hepatocytes with cell-type-specific induction of Cas9^16, 35^ (Materials and Methods). The library contained 456 sgRNAs targeting 202 genes, with most genes represented by two independent sgRNAs, as well as non-targeting controls. Each sgRNA was associated with a 185-nt barcode. The targeted genes spanned diverse aspects of liver biology, including metabolism, signal transduction, protein synthesis, and intracellular trafficking. We jointly measured the perturbation barcodes and 308 endogenous transcripts using MERFISH, and β-catenin to support cellular segmentation using oligo-conjugated antibody (Materials and Methods). The 308-gene endogenous panel was designed to capture multiple layers of liver biology.

**Fig. 2.**
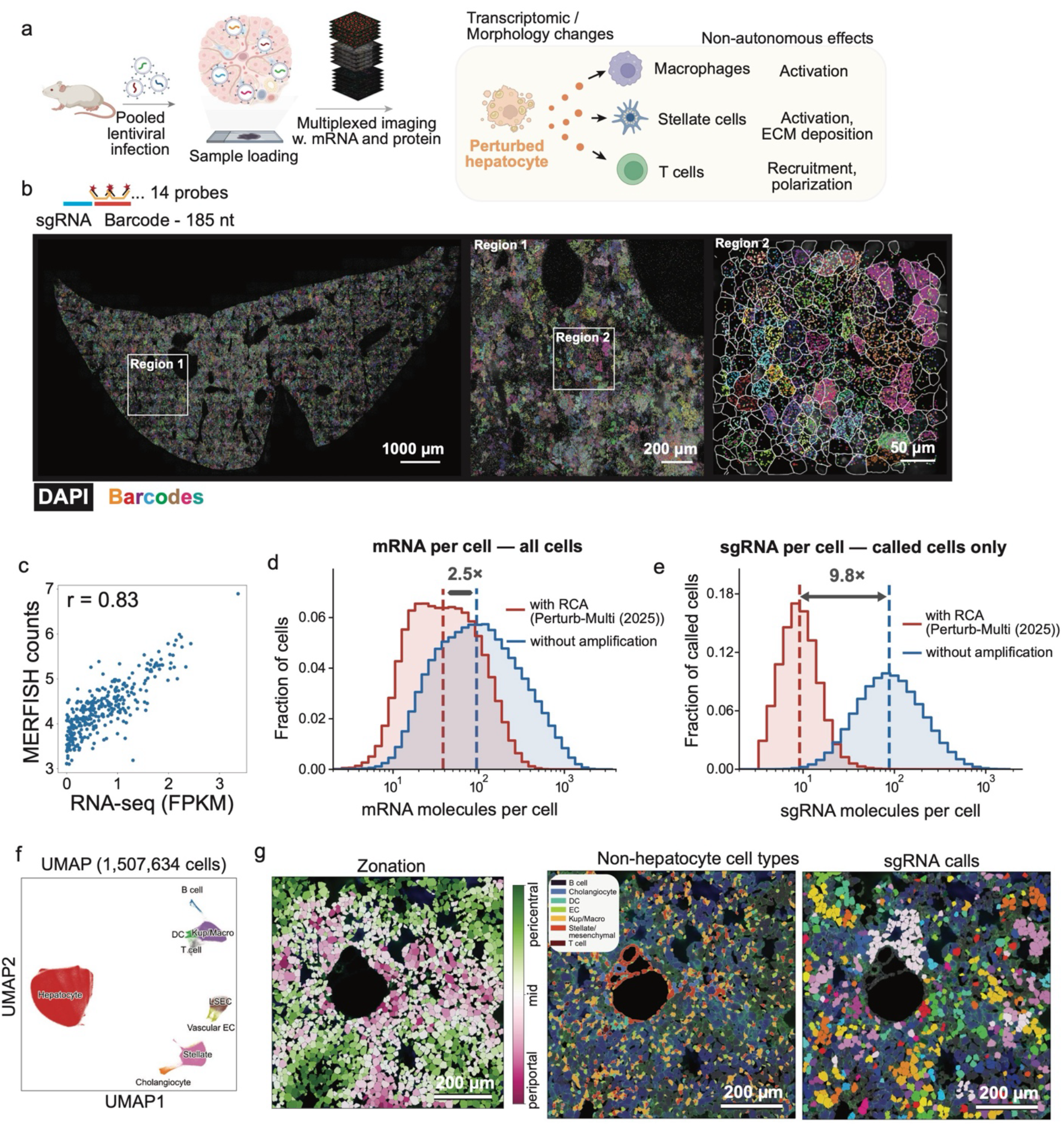
*In vivo* pooled genetic screen in mouse liver with amplification-free Perturb-MERFISH for sgRNA and mRNA detection. **a,** Schematic illustrating the application of amplification-free Perturb-MERFISH in mouse tissues to measure molecular and cellular phenotypes of genetic perturbations, including both cell-autonomous and non-autonomous effects. Some elements in this panel were created with BioRender. Jin, K. (2026) https://BioRender.com/yww91zb. **b,** Detection of sgRNA-associated barcodes in mouse liver using 185-nt barcodes. Distinct barcode identities, corresponding to different sgRNAs, are shown in different colors and overlaid with DAPI-stained nuclei. A global tissue mosaic is shown on the left, with two successive magnifications of the indicated regions in the middle and on the right. Segmented cell boundaries are also shown in the right image. **c,** Comparison of MERFISH mRNA counts with bulk RNA-seq measurements in wild-type mouse liver processed using the Perturb-MERFISH workflow. Each point represents a gene; Pearson correlation coefficient, *r* = 0.83. **d, e**, Comparison of endogenous mRNA (**d**) and sgRNA-barcode (**e**) detection performance in amplification-free Perturb-MERFISH with our previous results using RCA-based amplification^16^. Blue histogram: this work. Red histogram: our previous work based on RCA-MERFISH detection. **d**, Distribution of total mRNA counts per cell for the 103 genes shared between the targeted gene panels measured in these two studies. Dashed lines indicate median mRNA counts per cell, corresponding to a 2.5-fold increase with amplification-free Perturb-MERFISH. **e**, Distribution of sgRNA-barcode counts per cell for the 456-sgRNA library used in both studies. To enable a comparable assessment of sgRNA detection, only hepatocytes assigned a single perturbation were included from each dataset. Dashed lines indicate median sgRNA counts per cell, corresponding to a 9.8-fold increase with amplification-free Perturb-MERFISH. **f**, UMAP visualization of different cell populations detected in five liver samples classified using the transcription profiles of individual cells in the amplification-free Perturb-MERFISH. Cells are colored by annotated cell type. **g,** Representative spatial maps from a sub-region in Sample 1 showing hepatocyte zonation scores (left), annotated non-hepatocyte cell types (middle), and assigned sgRNA identities (right). Hepatocyte zonation scores are shown along the periportal-to-pericentral axis, calculated from the relative expression of selected pericentral and periportal marker genes (Materials and Methods), whereas non-hepatocyte cell types and sgRNA assignments are indicated by distinct colors. Scale bars are indicated in the images.

It contained markers for the major hepatic cell types (hepatocytes, Kupffer cells and other macrophages, dendritic cells, lymphocytes, liver sinusoidal and vascular endothelial cells, hepatic stellate cells, and cholangiocytes), as well as pericentral and periportal markers to resolve hepatocyte zonation. To measure perturbation responses, the panel further included genes that were differentially expressed in our previous Perturb-seq measurements using the same perturbation library^16^, covering stress-response, metabolic, and signaling pathways, together with ligands, receptors, and cell-state markers for non-parenchymal cells to enable detection of cell non-autonomous responses. Of the 308 genes, 103 overlapped with the panel used in our previous Perturb-Multi study^16^, enabling direct comparison of detection efficiency.

Direct hybridization without target amplification produced bright, cell-restricted barcode signals in liver tissue and assigned distinct sgRNA identities to individual cells throughout the tissue, revealing a broadly distributed mosaic of perturbations interspersed with local groups of cells (colonies) carrying the same sgRNA (Fig. 2b).

Across five tissue datasets, cell segmentation and quality-control filtering yielded 1,507,634 cells in total, including cells with and without assigned perturbation barcodes (Supplementary Figs. 2b–c, 3a). We first used DAPI and β-catenin staining to predict nuclear and cellular boundaries with a custom Cellpose model^36, 37^, then refined these boundaries using RNA distributions with Baysor^38^ (Materials and Methods). This combined approach recovered some cells missed by the initial Cellpose-based segmentation (Supplementary Figs. 2b–c). Following transcript-count and gene-count filtering, marker-based checks removed 14,358 putative segmentation doublets (0.9% of the remaining cells), including non-hepatocytes with excessive hepatocyte-marker counts and lymphocytes with excessive stellate-cell or endothelial-marker counts. Remaining cells contained an average of approximately 580 endogenous transcript counts across the 308-gene panel (Supplementary Fig. 3a). Aggregate transcript abundance per gene detected in this amplification-free Perturb-MERFISH protocol correlated well with bulk RNA-seq measurements, with a Pearson correlation coefficient of *r* = 0.83 (Fig. 2c).

### Direct amplification-free detection improves molecular recovery and perturbation assignment

Direct hybridization without target amplification increased molecular recovery relative to our previous RCA-MERFISH workflow. Among the 103 genes shared between the MERFISH endogenous gene panels, the median number of detected transcripts per cell increased from 38 to 95 in the current work, a 2.5-fold improvement (Fig. 2d). Across the 456-sgRNA library, the median number of barcode molecules corresponding to the assigned sgRNA per perturbation increased from 9 to 88 in singlet cells (cells with a single perturbation), a 9.8-fold improvement (Fig. 2d). These gains provide more molecular information both for perturbation assignment, increasing the number of informative cells, and for endogenous transcript profiling, increasing the sensitivity in perturbation response measurements. Avoiding sequential enzymatic reactions in RCA may reduce cumulative losses, although the contribution of individual processing steps was not measured.

Unsupervised analysis of endogenous transcript profiles resolved the major cell populations of the liver^39–41^ (Fig. 2f, Supplementary Fig. 3b). Hepatocytes formed the largest population and were readily distinguished from Kupffer cells and other macrophages^42, 43^, dendritic cells, B and T lymphocytes, liver sinusoidal endothelial cells (LSECs)^44, 45^, vascular endothelial cells, hepatic stellate cells^46^, and cholangiocytes (Fig. 2f, Supplementary Fig. 3b). Spatial projection of these annotations recapitulated the cellular organization of the intact tissue, with hepatocytes exhibiting a continuous periportal-to-pericentral gradient^39, 47, 48^ (Fig. 2g, Supplementary Fig. 3c) and non-hepatocyte populations distributed throughout the parenchyma and associated with vascular and biliary structures (Fig. 2g).

Across the five datasets, 425,487 hepatocytes received a single high-confidence perturbation call, yielding a median of 1,532 perturbed cells per targeted gene in these singlets (Supplementary Fig. 3d-e; Materials and Methods). A substantial fraction of hepatocytes carried multiple distinct perturbations (multiplets, 27.3%, approximately 257,000 cells), offering opportunities to investigate combinatorial perturbation effects in the future. Here, we focus on perturbation singlets in the subsequent analyses.

Mapping perturbation identities back into the tissue revealed small, spatially contiguous groups of cells carrying the same sgRNA. These groups are consistent with the clonal expansion of hepatocytes initially transduced during the neonatal period^16, 49^. The resulting perturbation territories were interspersed across the liver and sampled different positions along the lobular axis, enabling perturbation effects to be evaluated while retaining each cell’s local tissue context (Fig. 2g).

Together, these results show that amplification-free direct hybridization can jointly recover perturbation identity, endogenous gene expression, cell type, zonal state, and spatial context across a large number of cells in intact mouse liver.

### Widespread cell-autonomous transcriptional responses *in vivo*

We first examined cell-autonomous transcriptional responses in singly perturbed hepatocytes (Materials and Methods). Across the 202 targeted genes, the effect size varied, with many perturbations producing relatively few or no detectable changes in the expression of the 308 measured endogenous genes, while the strongest altered more than 100 genes (Fig. 3a). Strong phenotypes involved protein synthesis, proteostasis, intracellular trafficking, and hepatocyte identity. Transcriptional effects were reproducible across the five tissue samples, with pairwise correlations ranging from 0.77 to 0.87 (Supplementary Fig. 4a). We describe a few examples of cell-autonomous effects below for illustrative purposes, but focus on more systematic cell-non-autonomous effects in the following section.

**Fig. 3.**
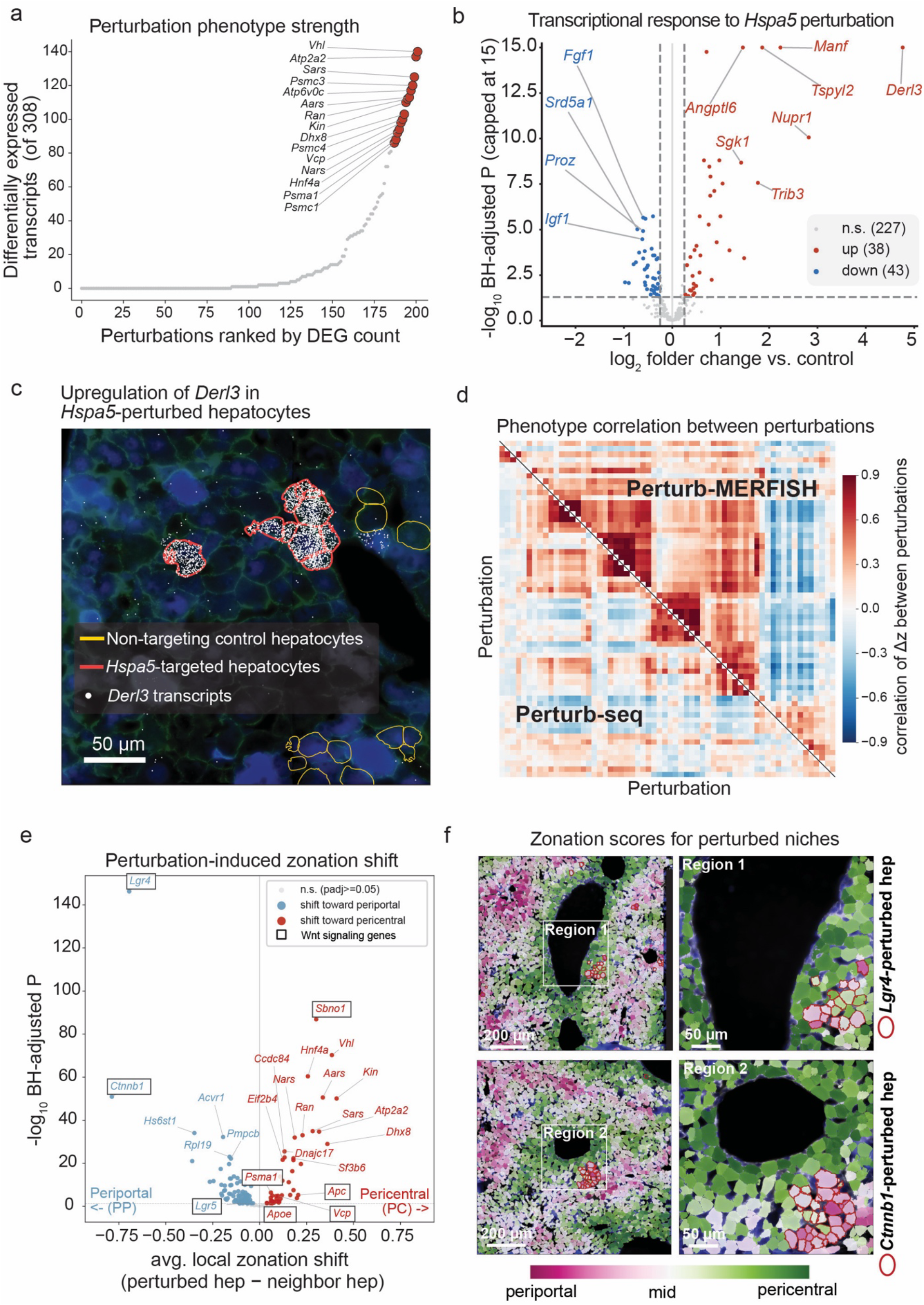
Perturb-MERFISH reveals widespread cell-autonomous effects of genetic perturbations in mouse liver. **a,** Number of differentially expressed genes (DEGs) detected for each of 202 perturbations, ranked by DEG count. Differential expression was tested among 308 measured endogenous mRNAs using per-sample pseudobulk limma-voom analysis of hepatocytes with a single perturbation assignment versus hepatocytes containing non-targeting sgRNA controls. Selected perturbations with strong transcriptional phenotypes are labeled. **b**, Volcano plot showing the transcriptional response to *Hspa5* perturbation relative to non-targeting controls. The y axis shows the negative log10-transformed Benjamini– Hochberg (BH)-adjusted *P* value, capped at 15. Significantly upregulated and downregulated genes are shown in red and blue, respectively; selected genes are labeled. Dashed lines indicate the statistical significance and effect-size thresholds used to identify DEGs. **c,** Representative Perturb-MERFISH image showing the drastic upregulation of *Derl3* transcription within *Hspa5*-targeted hepatocytes (outlined in red) vs non-targeting controls (outlined in yellow). *Derl3* transcripts are shown as white puncta, with DAPI-stained nuclei in blue and membrane staining (antibody against β-catenin) in green. Scale bar, 50 μm. **d,** Pairwise correlation matrix of perturbation-response profiles measured by Perturb-MERFISH and fixed-cell Perturb-seq for 62 perturbations with significant phenotypes in both datasets. Each entry represents the correlation between z-scored expression-change profiles for a pair of perturbations. Perturbations are shown in a shared consensus ordering; the upper and lower triangles display Perturb-MERFISH and Perturb-seq measurements, respectively. **e,** Perturbation-induced changes in hepatocyte zonation score relative to the unperturbed hepatocyte in the local tissue environment. For each perturbed hepatocyte, the zonation score was compared with the mean score of its ten nearest background unperturbed hepatocyte neighbors, and the mean difference for each perturbation is plotted against its BH-adjusted significance. Negative and positive values indicate a relative shift toward a periportal and pericentral state, respectively. Significant periportal and pericentral shifts are shown in blue and red, respectively; nonsignificant perturbations are shown in gray. Significant perturbations targeting Wnt pathway genes were highlighted with black boxes. Wnt pathway genes were defined as those annotated to either Reactome R-HSA-201681 (TCF-dependent signaling in response to WNT) or GO:0060070 (canonical Wnt signaling pathway). **f,** Representative spatial images showing clusters of *Lgr4*- or *Ctnnb1*-perturbed hepatocytes within their local tissue niches. For each perturbation, a larger sample region is shown on the left and a magnified view of the indicated region on the right. Hepatocytes are colored by zonation score from periportal to pericentral, and *Lgr4*- or *Ctnnb1*-perturbed hepatocytes are outlined in red. Both *Lgr4* and *Ctnnb1* perturbations are associated with a periportal shift in zonation score relative to neighboring hepatocytes within the same local tissue region.

*Hspa5*, which encodes the endoplasmic reticulum chaperone BiP^50, 51^, provides a direct illustration of these cell-autonomous responses. *Hspa5* disruption significantly upregulated 38 genes and downregulated 43 genes relative to controls (Fig. 3b). Upregulated genes included stress-response components *Derl3*, *Manf*, *Trib3*, *Nupr1*, and *Sgk1*^3, 52–57^, whereas hepatocyte functional genes including *Fgf1*, *Srd5a1*, *Proz*, and *Igf1* were reduced. Visually, *Derl3* transcripts showed a drastic upregulation in expression within *Hspa5*-perturbed hepatocytes, while control hepatocytes expressing non-targeting sgRNAs and most surrounding hepatocytes contained few *Derl3* transcripts, showing that the response was spatially restricted to perturbed cells (Fig. 3c, Supplementary Fig. 4b). We observed a similar trend for cells with the sgRNA targeting the ERAD factor *Sel1l*^58, 59^ (Supplementary Fig. 4b).

We next compared the cell-autonomous responses detected by Perturb-MERFISH with fixed-cell Perturb-seq measurements obtained using the same perturbation library^16^. Across perturbations with detectable phenotypes, the two modalities showed broadly similar transcriptional responses (median r = 0.7 for changes significant in at least one modality) and comparable maps of perturbation-perturbation response correlation (Fig. 3d), nominating functional modules of genes showing similar transcriptional response.

Spatial measurements also recovered expected changes in hepatocyte zonal identity, consistent with our previous Perturb-Multi study^16^. For example, comparing each perturbed hepatocyte with nearby background hepatocytes at a similar anatomical position, we found that *Lgr4* or *Ctnnb1* disruption shifted expression toward a periportal zonal identity, whereas *Apc* disruption shifted it toward a pericentral zonal identity^60–63^ (Fig. 3e–f, Supplementary Fig. 4c; Materials and Methods). Spatial visualization confirmed that these effects did not arise from preferential localization of perturbed cells: contiguous groups of *Lgr4*- or *Ctnnb1*-perturbed hepatocytes displayed markedly more periportal expression states than immediately surrounding hepatocytes occupying the same tissue region (Fig. 3f). These results directly demonstrate that Wnt signaling maintains hepatocyte zonal identity *in situ*^16, 60, 62, 63^ and illustrate how spatial perturbation screening can separate transcriptional state from physical position. *Vhl* disruption also induced a more pericentral-like state, consistent with activation of hypoxia-response programs that characterize the relatively low-oxygen pericentral environment, as observed previously^16, 39^. We also note that broadly acting perturbations can change the zonation score through general transcriptional remodeling, so the change in the summary zonation score as shown in Fig. 3e does not necessarily by itself establish selective regulation of zonal identity.

### Systematic mapping of cell non-autonomous perturbation phenotypes

We next mapped two complementary types of cell non-autonomous response: changes in local cell-type composition and changes in the transcriptional states of individual cell populations neighboring perturbed hepatocytes.

#### Genetic perturbations remodel local niche composition

For each hepatocyte carrying a single perturbation, we tested whether non-hepatocyte populations were enriched or depleted within 50 µm relative to control hepatocytes expressing non-targeting sgRNAs (Fig. 4a; Materials and Methods). We required a significant effect in the same direction in at least three of five liver samples. Many hepatocyte perturbations altered the relative abundance of neighboring vascular and sinusoidal endothelial cells, stellate cells, Kupffer cells/macrophages, dendritic cells, lymphocytes, and cholangiocytes (Fig. 4b). Distinct perturbations were associated with selective changes in particular cell-type populations, while some affected multiple populations. For example, *Vhl* perturbation was associated with enrichment of both LSECs and B cells, and perturbations affecting proteostasis (*Psma1*, *Psmc3*) and trafficking (*Ran*, *Copb2*, *Sec61g*) were frequently associated with increased Kupffer-cell/macrophage representation. These results motivated closer examination of selected local niche phenotypes.

**Fig. 4.**
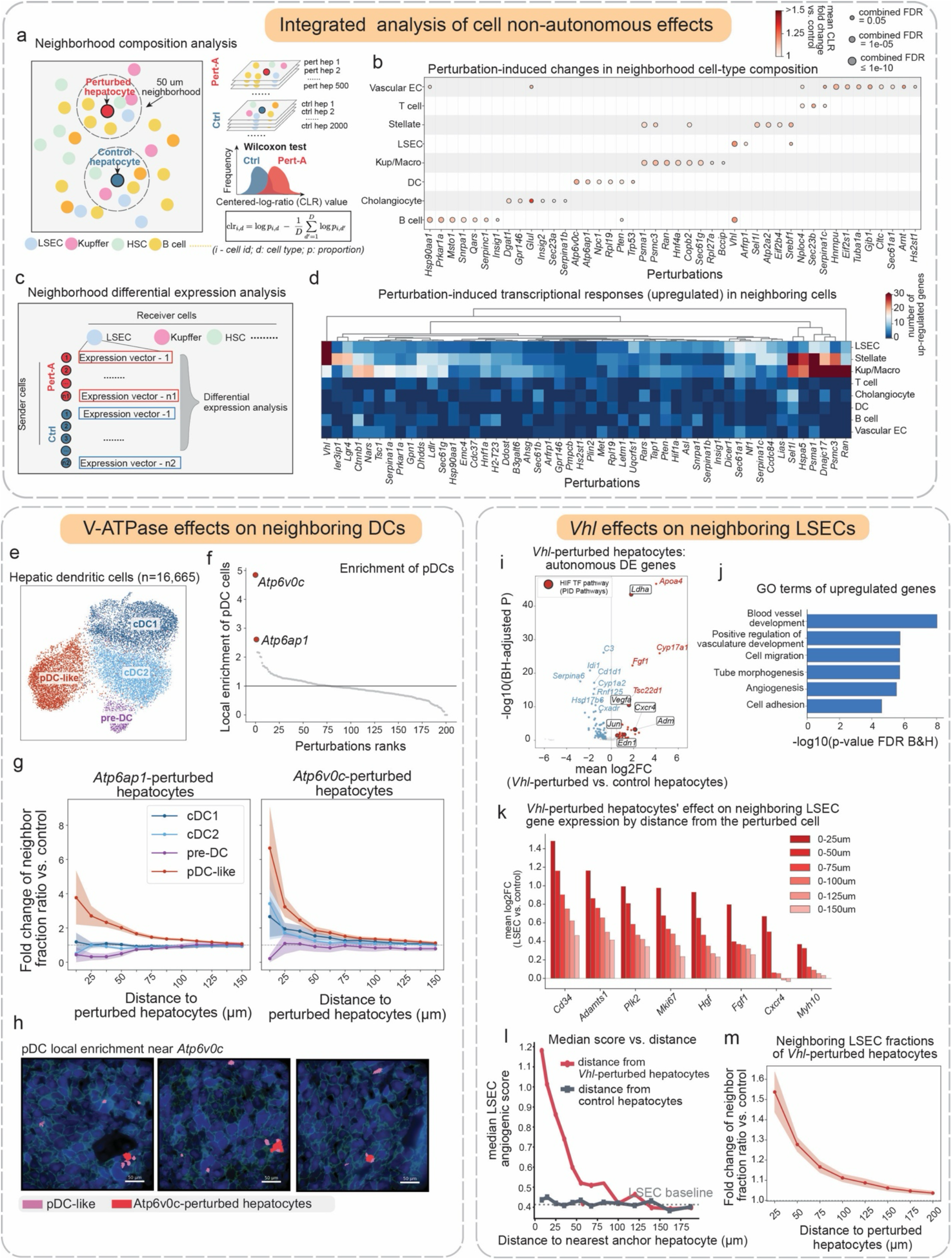
Perturb-MERFISH maps perturbation-induced cell-non-autonomous effects and remodeling of the liver microenvironment. **a,** Schematic of neighborhood cell-type composition analysis. For each hepatocyte carrying a single high-confidence perturbation, neighboring non-hepatocyte cells within a 50-μm radius were used to define its local niche, with analogous niches constructed around control hepatocytes. The relative abundance of each neighboring cell type was quantified for each niche and transformed using the centered log-ratio (CLR). For each cell type, CLR values across perturbed and control niches were compared using a Wilcoxon rank-sum test to identify perturbation-associated changes in local cellular composition (Materials and Methods). **b,** Perturbation-dependent changes in the cellular composition of local hepatocyte neighborhoods. Rows denote neighboring non-hepatocyte cell types, and columns denote perturbations associated with at least one consistently enriched cell population. Associations between perturbation and neighboring cell type enrichment were considered consistent when significant enrichment relative to control was observed in at least 3 of 5 independent datasets using the statistical framework described in **a**. Dot color represents the mean fold change in neighborhood CLR value relative to control, whereas point size represents the combined false-discovery rate (FDR; Materials and Methods). **c,** Schematic of the neighborhood differential-expression analysis. For each sender-hepatocyte perturbation and receiver-cell type, expression profiles of receiver cells neighboring perturbed hepatocytes were compared with those of the same cell type neighboring control hepatocytes across tissue samples. **d,** Heatmap of transcriptional responses in neighboring cells. Heatmap entries show the number of significantly upregulated genes for each sender-perturbation–receiver-cell-type combination; perturbations are ordered by hierarchical clustering. **e,** UMAP representation of 16,665 hepatic dendritic cells, resolving cDC1 (5,487 cells), cDC2 (5,943 cells), pre-DC (900 cells), and pDC-like (4,335 cells) populations. **f,** Local enrichment score for pDC-like cells near perturbed hepatocytes for all 202 perturbations. Local enrichment score is defined in Materials and Methods. Perturbations are ranked by enrichment, with *Atp6v0c* and *Atp6ap1* highlighted. The horizontal line indicates no enrichment relative to control. **g**, Distance-dependent enrichment of dendritic cell subtypes surrounding *Atp6ap1*-perturbed (left) and *Atp6v0c*-perturbed (right) hepatocytes. The fraction of neighboring cells assigned to each dendritic cell subtype was calculated within the specified distance ranging from 25 to 150 µm around perturbed hepatocytes and compared with that around non-targeting-control hepatocytes. Points show the mean fold change in neighboring cell fraction across liver samples, and shaded regions indicate mean ± standard error of the mean (SEM). **h,** Representative tissue regions showing local accumulation of pDC-like cells (magenta) near *Atp6v0c*-perturbed hepatocytes (red), overlaid on the DAPI-stained nuclei (blue) and membrane staining (green). Scale bars, 50 µm. **i,** Cell-autonomous transcriptional response of *Vhl*-targeted hepatocytes. Each point represents one measured gene. The x-axis shows the mean log2 fold change relative to control hepatocytes across samples; the y-axis shows the negative log10-transformed mean Benjamini–Hochberg-adjusted *P* value across samples. Genes with directionally consistent increases or decreases across samples are shown in red or blue, respectively; genes without consistent differential expression are shown in gray. Black outlines indicate genes in the HIF transcription factor pathway from the Pathway Interaction Database (PID). Selected genes are labeled. **j,** Gene Ontology Biological Process terms enriched among genes upregulated in LSECs neighboring *Vhl*-targeted hepatocytes. Bars show the negative log10-transformed BH-adjusted enrichment *p* value. **k,** Mean expression changes of selected genes in LSECs surrounding *Vhl*-targeted hepatocytes relative to LSECs surrounding control hepatocytes. Bars show mean log2 fold changes calculated using neighborhoods of different distance ranges from the anchor hepatocyte. **l,** Median angiogenic-response score of LSECs with successive distance bins from the nearest *Vhl*-targeted or control hepatocyte. The dashed horizontal line indicates the background LSEC score. **m**, Distance-dependent enrichment of LSECs surrounding *Vhl*-targeted hepatocytes. The fraction of neighboring cells classified as LSECs was calculated within the specified distance, ranging from 25 to 200 µm, around *Vhl*-targeted hepatocytes and compared with that around non-targeting-control hepatocytes. Points indicate the mean fold change in neighboring LSEC fraction across liver samples, and shaded regions indicate the SEM.

#### V-ATPase perturbation in hepatocytes is associated with localized pDC-like cell accumulation

One of the strongest immune-cell phenotypes was an enrichment of dendritic cells near hepatocytes targeted for *Atp6v0c*, which encodes a core membrane subunit of the V-ATPase^64, 65^. We therefore asked whether this screen-level phenotype reflected a general increase in dendritic cells or a response by a particular dendritic-cell state.

Subclustering the dendritic-cell/monocyte–macrophage compartment using MERFISH expression profiles identified four dendritic-cell populations comprising 16,665 cells, annotated as cDC1, cDC2, pre-DC, and pDC-like cells^66–69^ (Fig. 4e). We use the conservative term pDC-like because the panel was designed for broad liver-cell annotation rather than comprehensive dendritic-cell classification (Supplementary Fig. 5a). The *Atp6v0c*-associated phenotype was selective for this population: pDC-like cells were enriched 4.8-fold within 0–15 µm of perturbed hepatocyte centroids relative to non-targeting controls (Fig. 4f), whereas the other dendritic-cell populations showed weaker enrichment in the broader neighborhood analysis (Fig. 4g). pDC-like cell enrichment declined with distance and approached baseline in the 100–150-µm annulus (Materials and Methods). Two-dimensional density maps and tissue images showed pDC-like cells concentrated immediately around *Atp6v0c*-targeted hepatocytes, including cells with adjoining segmentation boundaries (Fig. 4h, Supplementary Fig. 5b).

We next quantified pDC-like cell enrichment within 0–15 µm of perturbed hepatocyte centroids across all 202 targeted genes (Materials and Methods). *Atp6v0c* produced the strongest phenotype, followed by *Atp6ap1*, which encodes an accessory V-ATPase component^65, 70^ (Fig. 4f). *Atp6ap1* perturbation produced 2.6-fold enrichment in this annulus. Both perturbations met the significance, effect-size, spatial-decay, and cross-section reproducibility criteria (Materials and Methods), with enrichment observed in all five sections. Both also showed enrichment using the separate, broader distance-dependent neighborhood metric (Fig. 4g). Their convergence supports an association between impaired hepatocyte V-ATPase function and localized pDC-like cell accumulation.

Lysosomal inhibition can increase extracellular-vesicle release and promote secretion of mitochondrial material, and stressed hepatocytes can release immunomodulatory extracellular vesicles^71, 72^. These findings suggest altered release of cellular material as one possible link between hepatocyte V-ATPase dysfunction and pDC-like cell accumulation. The localized enrichment could reflect recruitment, retention, survival, or another response of pDC-like cells, which awaits future experiments to discriminate.

#### Genetic perturbations alter transcriptional states of neighboring cells

Beyond changing local cell-type composition, perturbed hepatocytes could alter the molecular states of neighboring cells. We therefore compared expression in each neighboring non-hepatocyte population near gene-targeted hepatocytes with expression in the same population near control hepatocytes expressing non-targeting sgRNAs, testing for reproducible responses across samples (Fig. 4c; Materials and Methods).

Applying this analysis across the major non-hepatocyte populations revealed widespread cell non-autonomous transcriptional responses (Fig. 4d). The number and magnitude of responses varied substantially across both sender perturbations and receiver-cell types. LSECs, stellate cells, and Kupffer cells/macrophages exhibited the broadest responses, whereas lymphocytes and cholangiocytes showed fewer detectable changes. Individual perturbations could affect multiple receiver cell populations or act selectively on particular cell types. *Vhl* was among the most pleiotropic perturbations, producing responses in multiple receiver cell populations, with the effects being particularly strong on LSECs and stellate cells, while also having a substantial response in Kupffer cells. We therefore focused on the effects of the hepatocyte *Vhl* perturbation on LSEC and stellate cells below.

#### Vhl disruption induces localized endothelial responses

*Vhl* disruption produced the strongest transcriptional phenotype in neighboring LSECs (Fig. 4d, Supplementary Fig. 5c). *Vhl*-perturbed hepatocytes activated a cell-autonomous hypoxia-response program^73, 74^, including induction of angiogenic factors such as *Vegfa* and *Adm*^75, 76^ (Fig. 4i). These changes suggest HIF-associated signaling as a candidate mechanism for the neighboring endothelial response.

LSECs near *Vhl*-perturbed hepatocytes exhibited a transcriptional response enriched for genes associated with blood-vessel development, angiogenesis, cell migration, tube morphogenesis, and cell adhesion (Fig. 4j). Upregulated genes included *Cd34*, *Adamts1*, *Plk2*, *Mki67*, *Hgf*, *Fgf1*, and *Cxcr4* (Fig. 4k). Expression was highest near *Vhl*-perturbed hepatocytes and declined with increasing distance (Fig. 4k). A composite angiogenic-response score likewise fell sharply over approximately the first 50 µm and approached background at greater distances (Fig. 4l).

*Vhl* disruption also altered the local abundance of LSECs. LSECs constituted a larger fraction of cells immediately surrounding *Vhl*-perturbed hepatocytes than around control hepatocytes expressing non-targeting sgRNAs, with the difference diminishing as distance from the perturbed hepatocyte increased (Fig. 4m, Supplementary Fig. 5d).

Hepatic stellate cells (HSCs) near *Vhl*-perturbed hepatocytes also showed pronounced transcriptional changes (Fig. 4d), including increased expression of contractile and activation-associated genes *Acta2, Myl9, Tpm2,* and *Tagln*, and the extracellular-matrix gene *Col3a1*^46, 77^ (Supplementary Fig. 5e). These changes were strongest immediately adjacent to *Vhl*-perturbed hepatocytes and declined with distance, consistent with localized HSC activation (Supplementary Fig. 5f).

Previous hepatocyte-directed *Vhl* inactivation in mice caused increased angiogenic gene expression and vascular remodeling, including endothelial proliferation, with subsequent genetic studies demonstrating a major role for hepatocyte Hif2α (*Epas1*) in this response^78, 79^. Here, we demonstrate how spatial screens can turn this physiological response into a quantitative readout by identifying the responding receiver-cell populations, defining their induced transcriptional programs, and measuring how those responses vary with distance from *Vhl*-perturbed hepatocytes.

## Discussion

Here we developed a simplified and yet more efficient and scalable *in situ* perturbation detection method for imaging-based pooled genetic screening. Using this amplification-free Perturb-MERFISH approach, the detection efficiencies of perturbation barcodes and endogenous mRNA transcripts increased by 9.8-fold and 2.5-fold, respectively, compared with our previous RCA-based workflow^16^. These gains show that amplification is not required for reliable assignment of highly expressed perturbation barcodes, suggesting that losses during amplification, potentially because of the limited efficiency and variability of enzymatic reactions, can outweigh its resulting signal gain on some transcripts. The increased perturbation barcode and endogenous mRNA counts per cell improved the efficiency of perturbation assignment and the detection sensitivity to cellular responses, substantially increasing the number of informative cells per experiment and enabling robust analysis of not only cell-autonomous but also cell non-autonomous phenotypes. Our observation that barcodes as short as 90 nt could be reliably detected further supports compact designs that simplify pooled library construction.

Applying this amplification-free Perturb-MERFISH to a mouse liver screen targeting 202 genes, we profiled more than 1.5 million cells, and identified many non-autonomous effects of perturbed hepatocytes on the abundance and transcriptional state of neighboring cells. For example, localized enrichment of pDC-like cells was observed near hepatocytes with disruption of *Atp6v0c* or *Atp6ap1*, components of the V-ATPase. *Vhl* disruption, which activated a hypoxia program in hepatocytes, induced angiogenic responses and enrichment of nearby LSECs, together with localized HSC activation. Follow-up studies with targeted and/or combinatorial perturbations of candidate signaling pathways, broader molecular profiling, and time-resolved imaging could determine whether the cell accumulation reflects recruitment, proliferation, retention, or survival, as well as the molecular mechanisms underlying the enrichment and molecular state changes of these cells.

By jointly measuring perturbation identity, cell state, and spatial context, Perturb-MERFISH provides a scalable way to investigate how genetically altered cells reshape both the perturbed cells themselves, and their neighboring cells and local microenvironment, in intact tissues. Currently, the targeted perturbation and endogenous RNA panels capture only a limited portion of the perturbation space and part of the cellular response to these perturbations. Future studies that increase the perturbation coverage and the numbers of endogenous RNAs and proteins measured will expand the breadth and depth of gene function identification. This approach could also be adapted to other perturbation modalities, including CRISPR interference^80, 81^, CRISPR activation^33, 82^, base editing^83^, and overexpression libraries^84^. Expanded phenotypic measurements and greater perturbation space coverage, including combinatorial perturbations^32, 85, 86^, will extend the range of molecular mechanisms of cell and tissue functions accessible to these screens.

## Acknowledgements

We thank T. Norman and C. Chu for helpful discussions. This work is in part supported by the Howard Hughes Medical Institute (HHMI). R.A.S. is a Junior Fellow at Harvard Society of Fellows. J.S.W. and X.Z. are HHMI investigators. AI models from OpenAI and Anthropic assisted with exploratory data analysis and manuscript editing.

## Competing interests

K.J., R.A.S., W.E.A., J.S.W. and X.Z. are inventors on patents applied for by Harvard University and Whitehead Institute related to imaging-based screening. R.A.S. and W.E.A. consult for Xaira Therapeutics. J.S.W. declares outside interest in 5 AM Venture, Amgen, nChroma Bio, DEM Biosciences, KSQ Therapeutics, Maze Therapeutics, Tenaya Therapeutics, Tessera Therapeutics, Thermo Fisher Scientific, and Xaira Therapeutics. X.Z. is a co-founder and consultant of Vizgen, Inc.

## Materials and Methods

### Animals

Animal husbandry, viral delivery, tissue collection, and sectioning were performed as described previously^16^. Experiments used male and female wild-type mice and B6;129-Gt(ROSA)26Sor^tm1(CAG-Cas9*,-EGFP)Fezh^/J mice maintained on a C57BL/6J background^35^. Animals were purchased from The Jackson Laboratory or bred at the Massachusetts Institute of Technology.

Mice were housed at 22 ± 1°C and 30–70% relative humidity under a 12-hour light/dark schedule, with the dark phase extending from 14:00 to 02:00. Food and water were available without restriction except during fasting experiments. Animals received standard chow (Research Diets, D12492i).

Lentivirus was administered at postnatal day 1, followed by hepatocyte-directed AAV-Cre delivery at approximately postnatal day 30. Tissues were collected 10 days after AAV administration.

All procedures conformed to NIH guidelines and were approved by the Committee on Animal Care at MIT.

### Cell culture

Cell-culture procedures followed those reported previously^16^. HEK 293T/17 cells (ATCC CRL-11268) were maintained in DMEM containing GlutaMAX, HEPES, 10% fetal bovine serum, penicillin, and streptomycin.

AML12 cells (ATCC CRL-2254) were cultured in DMEM/F-12 containing 10% fetal bovine serum, insulin, transferrin, selenium, and dexamethasone. Cell-culture media and supplements were obtained from Thermo Fisher Scientific.

CRISPRi RPE1 cells were described previously^6^ and were grown in DMEM/F-12 supplemented with 10% FBS, 100 units/mL penicillin, and 100 μg/mL streptomycin.

### CRISPR library construction

Guide-barcode libraries for the *in vitro* experiments and the 185-nt *in vivo* experiments were generated as described previously^16^. Guide-barcode libraries with 260-nt barcodes were generated in a two-step procedure. First, an sgRNA expression vector with a U6 promoter and no sgRNA constant region was digested with BstXI and BlpI and an amplified oligonucleotide library (Twist Bioscience) with an sgRNA targeting region, two BsmBI cut sites, and a perturbation barcode was inserted by ligation. The library was then digested with BsmBI and a pool of three sgRNA constant regions^3^ (CR1, CR2, CR3) was inserted by Gibson assembly. The use of three constant regions minimizes recombination between the sgRNA targeting region and the perturbation barcode during lentiviral packaging and transduction.

### Lentivirus production

Lentiviral libraries were produced in HEK 293T/17 cells using standard packaging procedures described previously^16^. Cells were transfected with the library plasmid, psPAX2, and pMD2.G using FuGENE HD. ViralBoost was added according to the manufacturer’s instructions.

For *in vivo* experiments, viral supernatant was collected, passed through a 0.45-µm PES filter, and concentrated approximately tenfold by polyethylene glycol precipitation using Lenti-X Concentrator. The resulting material was further concentrated by centrifugation at 25,000 × g for 2 hours at 4°C in an SW 32 Ti rotor. Viral pellets were resuspended in cold PBS containing 4% glucose, flash frozen, and stored until use. Functional titers were estimated by transduction of AML12 cells.

For *in vitro* experiments, viral supernatant was collected and clarified by centrifugation at 500 × g for 5 min at 4°C to remove residual cells and cellular debris. The clarified supernatant was subsequently mixed with Lenti-X Concentrator at a 3:1 (supernatant: concentrator) volume ratio and incubated overnight at 4°C. Viral particles were pelleted by centrifugation at 1,500 × g for 45 min at 4°C, after which the supernatant was carefully discarded and the pellet was resuspended in one-tenth of the original supernatant volume, resulting in an approximately tenfold-concentrated viral preparation.

### Generation of mosaic livers

Mosaic livers were generated as described previously^16^. Concentrated lentivirus was thawed on ice, and up to 50 µL was delivered through the temporal vein of postnatal day 1 mice.

After the mice reached ∼30 days, Cas9 activity was induced by retro-orbital administration of approximately 5 × 10^11^ genome copies of AAV8 encoding Cre under a hepatocyte-selective promoter (Addgene 107787-AAV8). Animals were analyzed 10 days later.

Animals were anesthetized and perfused sequentially with PBS and 4% paraformaldehyde. Livers were removed, fixed for an additional 3 hours in 4% paraformaldehyde, and cryoprotected for approximately 16 hours in 4% paraformaldehyde containing 30% sucrose. Samples were embedded in optimal cutting temperature compound, frozen, and stored at −80°C.

### Backbone construct for barcode tests in RPE1 cells

The backbone mosaic screening vector was derived from a modified CROP-seq vector^4^ (pBA950; Addgene plasmid #122239; Replogle et al., 2020). The vector was further engineered by introducing a multiple cloning site (MCS) and a puromycin-resistance cassette, yielding the final backbone used to construct sgRNA libraries with different barcode lengths for testing in RPE1 cells.

### Design of sgRNA barcodes

To generate a pool of sgRNA barcodes, we first generated random sequences of target lengths. Candidate barcode sequences were computationally screened to minimize sequence features that could affect the expression, processing, stability, or detection of barcode-containing transcripts. Sequences were designed to have a GC content between 40% and 60%. Each candidate was screened for canonical and variant polyadenylation signals, AU-rich and GU-rich elements, extended poly(A) tracts, homopolymer runs, potential microRNA target sites, cytoplasmic polyadenylation and Pumilio-binding elements, and putative cryptic splice-donor, splice-acceptor, and branch-point motifs. Candidate sequences were additionally evaluated for direct sequence repeats that could increase the risk of recombination or synthesis artifacts. Detected features were classified as high-, medium-, or low-severity sequence liabilities, and candidates containing any high-severity feature were excluded. RNA secondary-structure potential was assessed using sequence-composition and reverse-complementarity heuristics, with optional minimum-free-energy prediction using RNAfold^87^. Barcodes passing these filters were selected for incorporation into the sgRNA–barcode constructs.

We further screened candidate barcode sequences to minimize complementarity to endogenous transcripts, PCR primer sequences, and MERFISH readout sequences, thereby providing orthogonal targets for MERFISH encoding probes. Specifically, the large pool of randomly generated barcode sequences described above was processed through the same MERFISH probe-design pipeline used in our previous studies^16, 30, 88, 89^. Candidate barcodes were ranked according to the number of valid MERFISH encoding probes that could be designed for each sequence after applying these specificity constraints. The number of designable probes therefore served as a proxy for sequence orthogonality to the endogenous transcriptome and other assay oligonucleotides. Barcode sequences with the largest numbers of designable MERFISH probes were selected for subsequent perturbation experiments.

### Design and preparation of MERFISH encoding probes

Encoding probes were designed separately for endogenous mRNAs and sgRNA barcodes. The mRNA probe library was generated using an established design workflow^89^ that evaluated all candidate 30-nt sequences complementary to the genes in our panel. Candidate sequences were screened for potential off-target matches against coding and noncoding transcripts and excluded if they showed substantial similarity to unintended targets. Additional selection criteria restricted GC content to 43–63% and predicted melting temperature to 66–76 °C^30, 88^. Sequences passing these filters served as the target-binding components of the encoding probes. Each encoding probe also incorporated 20-nt readout sequences, which provided binding sites for complementary fluorescent probes used in MERFISH imaging. Readout sequences used only A, T, and G nucleotides and followed previously reported design principles^88, 90^. Their design limited cross-reactivity and secondary structure to support efficient hybridization and reduce probe aggregation. The final oligonucleotide designs included primer-binding sites for library amplification^30^, with an average of 96 encoding probes per mRNA target.

For encoding probes targeting sgRNA barcodes, we applied design criteria similar to those used for the mRNA encoding probes. Because the barcode sequences were substantially shorter than mRNA transcripts, candidate probes were tiled at high density with extensive overlap, yielding approximately one encoding probe every 13–15 nt. This design produced approximately seven encoding probes for a 90-nt barcode and eleven encoding probes for a 150-nt barcode. To further enhance signal intensity, each encoding probe contained four to five readout sequences, enabling recruitment of multiple fluorescent readout probes during MERFISH imaging.

The encoding-probe libraries were synthesized as oligo pools by Twist Bioscience and amplified as previously described^89^. Briefly, the libraries were first amplified by qPCR to generate double-stranded DNA templates containing a T7 promoter. These templates were then subjected to overnight *in vitro* transcription to produce RNA intermediates. The RNA products were purified using Zeba desalting columns and reverse-transcribed to generate RNA–DNA hybrids. Following reverse transcription, the RNA strands were removed by alkaline hydrolysis, and the remaining single-stranded DNA probes were purified using Zymo Research columns. After concentration, the final encoding-probe libraries typically reached 10–40 nM per probe.

### Sample preparation for direct-hybridization experiments

For RPE1 cells, samples were fixed in freshly prepared 4% paraformaldehyde (PFA) in 1× PBS for 15–20 min at room temperature. The fixative was then removed, and the cells were washed three times with 1× PBS before permeabilization with 0.5% Triton X-100 in 1× PBS for 20 min on a rocking platform. After three washes with 2× SSC, the cells were hybridized with sgRNA encoding probes at 2 nM per probe in hybridization buffer containing Stellaris RNA FISH Hybridization Buffer, 30% (v/v) formamide, 2% murine RNase inhibitor, 1 µM poly(A) probe, and a reporter-gene smFISH probe library at 2 nM per probe. The poly(A) probe contained a mixture of DNA and locked nucleic acid nucleotides (/5Acryd/TTGAGTGGATGGAGTGTAATT+TT+TT+TT+TT+TT+TT+TT+TT+TT+T), where “+T” denotes an LNA-modified thymidine and /5Acryd/ indicates a 5′ acrydite modification. Hybridization was carried out for 36–48 h at 37 °C in a humidified chamber. Following hybridization, samples were washed twice for 30 min each in 30% (v/v) formamide and 0.1% Triton X-100 in 2× SSC. The samples were then equilibrated twice for 5 min each in gel solution containing 4% (v/v) acrylamide/bis-acrylamide (19:1), 300 mM NaCl, 60 mM Tris-HCl (pH 8.0), and 0.2% (v/v) tetramethylethylenediamine (TEMED). Immediately before polymerization, ammonium persulfate (APS) was added to a final concentration of 0.2% (w/v). Coverslips were inverted onto approximately 60 µL of the gel mixture placed on a glass surface and incubated for 2 h at room temperature. The embedded samples were then digested in 3 mL of digestion buffer for 1–2 days at 37 °C^88^. The digestion buffer contained 2% (v/v) sodium dodecyl sulfate (SDS), 2× SSC, 0.5% Triton X-100, and proteinase K diluted 1:50 (v/v). After digestion, the samples were transferred to a clean dish and washed four times with 2× SSC for 30 min per wash. Samples were either imaged immediately or stored for up to 3 days in 2× SSC supplemented with murine RNase inhibitor at a 1:100 dilution.

For mouse liver sections, we modified the protocol to improve RNA preservation and support simultaneous hybridization of mRNA and sgRNA-barcode encoding-probe libraries. Following cryosectioning, tissue sections were air-dried at 47 °C for 8–10 min and fixed in 4% paraformaldehyde (PFA) in 1× PBS for 15–20 min. Samples were washed three times with 1× PBS and stored overnight in 70% ethanol at 4 °C; under these conditions, sections could be stored for several weeks. Before further processing, samples were washed three times with 1× PBS and subjected to heat-mediated decrosslinking in 1× Tris–EDTA buffer (pH 9.0) at 70 °C for 4 h, followed by three washes with 1× PBS. For membrane staining, sections were blocked for 1 h at room temperature with 50 µL of blocking buffer containing 20% (v/v) ultrapure BSA, 0.5% dextran sulfate, 0.5% Triton X-100, 50 mM NaCl, and 0.5 mg/mL sheared salmon sperm DNA in 1× PBS. Samples were then incubated overnight at 4 °C in a humidified chamber with 1–2 µg/mL oligonucleotide-conjugated β-catenin antibody diluted in blocking buffer. Oligonucleotide-conjugated antibodies were prepared as previously described^16^. After incubation, samples were washed twice with 1× PBS for 5–7 min per wash, post-fixed in 4% PFA in 1× PBS for 20 min, and washed three times with 1× PBS. To improve RNA retention during subsequent gel embedding and digestion, tissue RNA was modified with MelphaX to enable anchoring to the polyacrylamide matrix. Specifically, after post-fixation, samples were equilibrated in 20 mM MOPS buffer (pH 7.7) for 30 min at room temperature. MelphaX stock solution (2 mg/mL) was diluted to 1 mg/mL in 40 mM MOPS buffer (pH 7.7), and 50 µL was applied to each section. Samples were incubated overnight at 37 °C and then rinsed twice with 1× PBS for 5 min per wash. MelphaX stock was prepared as previously described^91^. Samples were then embedded in 4% polyacrylamide gel and digested using the same conditions described above for cultured cells. Following digestion and washing with 2× SSC, sections were hybridized with both mRNA and sgRNA-barcode encoding-probe libraries at 1 nM per probe and 2 nM per probe, respectively. Hybridization conditions were identical to those used for RPE1 cells. After hybridization, samples were washed twice for 30 min each in 30% (v/v) formamide and 0.1% Triton X-100 in 2× SSC. Samples were then either imaged immediately or stored at 4 °C for up to 1 week in 2× SSC with murine RNase inhibitor at a 1:100 dilution.

### MERFISH imaging

MERFISH imaging was performed using a custom-built epifluorescence microscope coupled to an automated fluidics system, as described previously^16, 30, 88^. Before imaging, samples were stained with DAPI at a 1:100,000 (v/v) dilution, a poly(T) readout probe, and readout probes corresponding to the first three bits of the MERFISH codebook. Staining for these initial three bits was performed manually on the bench using the same hybridization conditions applied during subsequent automated imaging rounds, described below.

All imaging buffers were prepared immediately before use, degassed for 30 min, and loaded onto the fluidics system. An adaptor-based readout strategy was used. The hybridization buffer consisted of 30% (v/v) formamide and 0.1% Triton X-100 in 2× SSC. The cleavage buffer contained 50 mM tris(2-carboxyethyl)phosphine hydrochloride (TCEP-HCl), 30% (v/v) formamide, 0.1% Triton X-100, and 50 nM per probe of blocker probes in 2× SSC. The blocker probes had the same sequences as the fluorescent readout probes but lacked fluorophore conjugation. The wash buffer had the same composition as the hybridization buffer. The imaging buffer contained 2× SSC, 10% (w/v) glucose, 60 mM Tris–HCl (pH 8.0), 1 mg/mL glucose oxidase, catalase diluted 1:170 (v/v), 1 mg/mL Trolox, 50 µM Trolox quinone, and murine RNase inhibitor diluted 1:1,000. Adaptor and fluorescent readout probes were diluted in hybridization buffer to 10 nM and 5 nM per probe, respectively.

Samples were mounted in a commercial flow chamber (FCS2, Bioptechs) fitted with a 0.75-mm-thick gasket (DIE F18524, Bioptechs), and the chamber was filled with imaging buffer. During the first imaging round, three fluorescence channels corresponding to the 750-, 650-, and 560-nm readout probes were acquired together with the poly(T) signal at 488 nm and DAPI and fiducial-bead signals at 408 nm. For each field of view, six z-planes were collected at 1.5-µm intervals.

After completion of the first imaging round, fluorophores were removed from the readout probes by incubation in cleavage buffer for 15 min, followed by washing with 3 mL wash buffer. For each subsequent round, samples were hybridized with the corresponding adaptor probes for 20 min and washed with 3 mL wash buffer. Fluorescent readout probes were then applied for 20 min, followed by a second wash with 3 mL wash buffer. Imaging buffer was introduced before image acquisition. In these later rounds, the 750-, 650-, and 560-nm channels were acquired across six z-planes, whereas the 408-nm fiducial-bead channel was acquired at a single z-plane near the coverslip surface. Each imaging round measured three MERFISH bits, and the codebooks used in this study contained 14–26 bits in total.

### Image data processing

MERFISH image processing was performed using the Python package MERlin (https://github.com/ZhuangLab/MERlin), as described previously^16, 92^. Images from successive imaging rounds were first registered using fiducial-bead signals acquired in every field of view (FOV). For each FOV, image stacks were processed by high-pass filtering for background removal, followed by ten iterations of Richardson–Lucy deconvolution^93, 94^ and subsequent low-pass filtering. Individual RNA molecules were then identified using MERlin’s pixel-based barcode-decoding algorithm. Decoded barcodes were filtered based on their estimated misidentification rate at a 5% threshold. For sgRNA-barcode decoding in some liver tissue samples, a threshold of 1% was applied to further remove background signals. mRNA and sgRNA-barcode images were registered to the same reference imaging round. Because the two signal classes differed in intensity and background characteristics, image filtering, deconvolution, and barcode decoding were performed separately for mRNA and sgRNA data.

### Cell segmentation

Cell segmentation was performed using Cellpose v3^36, 37^ and Baysor v0.8.2^38^. Initial cell boundaries were predicted with a custom Cellpose model trained on DAPI nuclear staining and β-catenin membrane staining. The resulting Cellpose masks, together with the spatial coordinates of decoded mRNA molecules, were then used as inputs for three-dimensional segmentation with Baysor. This combined approach incorporated morphological information from both nuclear and membrane signals while allowing transcript distributions to refine cell boundaries and recover cells with weak or incomplete DAPI or membrane staining that were missed by Cellpose alone. Baysor segmentation was performed using -m 20 and --prior-segmentation-confidence 0.5. sgRNA molecules were assigned to cells via a point-in-polygon test against the cell boundaries produced by the Baysor mRNA segmentation. Molecules not falling within any cell boundary were excluded from downstream analysis.

### Single-cell data analysis of MERFISH data

Single-cell analysis was conducted using the Python package Scanpy^95^. After segmentation, cells with fewer than 50 mRNA counts or fewer than 10 detected genes were removed. Raw mRNA counts were normalized to counts-per-10,000 (CP10K) and log1p-transformed. The top 275 highly variable genes (of the 308-gene panel) were then selected using Scanpy’s Seurat-flavor method^96^, computed per sample and combined across samples, and the dataset was subset to these genes prior to downstream analysis. The data were then scaled (clipped at a maximum value of 10) and principal component analysis (PCA) was performed, retaining 30 principal components. Batch effects across the five imaging samples were corrected using Harmony^97^ applied to the principal component space. A neighborhood graph was constructed from the 30 Harmony-corrected principal components (40 nearest neighbors), followed by UMAP embedding^98,99^. Leiden clustering^100^ was performed using the igraph implementation at three resolutions: 0.5, 1.0, and 2.0. Cell types were annotated based on the expression of canonical marker genes enriched in each cluster, identified using Leiden clustering at a resolution of 2.0. For example, LSECs were identified by the expression of markers such as *Lyve1* and *Clec4g*.

Putative doublets arising from segmentation errors were removed using two sequential heuristic filtering steps based on raw transcript counts from curated marker-gene panels. First, non-hepatocyte cells exhibiting implausibly high expression of hepatocyte markers were classified as hepatocyte-contaminated doublets. Cells with a summed raw count greater than 20 across a panel of 21 hepatocyte-enriched genes—comprising 13 high-confidence markers, including *Apoa4*, *Cyp2c38*, and *Serpina7*, and eight supporting markers—were excluded. This step removed 9,220 of 1,521,992 cells (0.6%). Second, among cells annotated as T or B lymphocytes, cells showing substantial contamination from hepatic stellate cell or liver sinusoidal endothelial cell (LSEC) transcripts were removed. A lymphocyte was classified as a doublet if its summed raw count exceeded 20 across either a 10-gene stellate cell marker panel, including *Col1a1*, *Col1a2*, and *Pdgfrb*, or a 10-gene LSEC marker panel, including *Stab2*, *Pecam1*, and *Cdh5*. Together, these filtering steps reduced the dataset from 1,521,992 to 1,507,634 cells, corresponding to the removal of 14,358 cells (0.9%).

### Specificity test of perturbation calling

To assess the specificity of barcode detection, RPE1 cells were separately infected with libraries containing 90-, 150-, or 240-nt barcodes, and each sample was decoded against the combined codebook containing 77 barcode identities, including 41 blank barcodes. Each barcode-length library contained 12 distinct barcodes.

Misclassification to another barcode within the same barcode-length library cannot be directly distinguished from a correct assignment in this experiment. Assuming that erroneous barcode assignments are equally likely among incorrect barcode identities, the expected ratio of within-library to outside-library misclassifications is ((12-1)/(77-12)=0.169). Based on the 1 observed outside-library misclassifications, we therefore estimated (1 * 0.169 = 0.169) additional within-library misclassifications. The overall barcode-calling error rate was thus estimated to be ((1+0.169)/4,475 = 0.026%).

### sgRNA calling

Per-cell CRISPR perturbation identity was inferred from MERFISH-detected sgRNA barcode counts using a two-stage calling procedure. In the first stage, each sgRNA guide was called independently within each tissue sample, as samples were imaged separately and exhibited distinct background distributions. For a given guide, nonzero raw counts across cells were log10(count + 1)-transformed and modeled as a two-component Gaussian mixture with a shared (tied) covariance. The mixture component with the higher mean was designated the true-signal component, and cells with a posterior probability greater than 0.5 of belonging to this component were assigned the guide, with the detection threshold set to the minimum raw count among these cells. Guides with fewer than five nonzero-count cells or a maximum count below three were excluded from calling. A second stage applied a per-cell dominance criterion, retaining a guide assignment only if its count was at least 70% of that cell’s maximum guide count.

### Zonation score

For each hepatocyte, a pericentral (PC) module score and a periportal (PP) module score were computed as the mean log-normalized expression across a curated panel of eight PC marker genes (*Cyp2c38*, *Serpina7*, *Aldh3a2*, *Axin2*, *Pdk4*, *Lpl*, *Serpina6*, *Cyp1a2*) and eight PP marker genes (*Hsd17b6*, *Aldh1b1*, *Clec2h*, *Cib3*, *Cyp2b10*, *Car14*, *Igf1*, *Egfr*), respectively. A single zonation score per cell was then defined as PC_minus_PP = PC score − PP score, such that positive values indicate a more pericentral transcriptional identity and negative values a more periportal identity. Zonation scores were computed only for hepatocytes.

### Hepatocyte zonal shift analysis

To determine whether individual genetic perturbations shifted hepatocytes along the pericentral–periportal (PC–PP) zonation axis, we performed a local, spatially matched, neighbor-relative analysis. For each perturbed hepatocyte assigned a single sgRNA, we identified its *K* nearest eligible background hepatocytes within the same tissue sample using three-dimensional Euclidean distance in *x*, *y*, and *z*. Analyses were performed independently with *K* = 10, 20, and 30 to assess robustness. For each perturbed cell, we calculated a local zonation shift, delta, as its PC-minus-PP zonation score minus the mean score of its *K* neighboring background hepatocytes. Delta values were then pooled across all hepatocytes carrying the same perturbation across the five samples. A two-sided one-sample Wilcoxon signed-rank test against zero was used to determine whether each perturbation induced a consistent shift relative to the local tissue environment, with delta > 0 indicating a shift toward a pericentral-like state and delta < 0 indicating a shift toward a periportal-like state. Perturbations represented by fewer than 30 valid cells were excluded. P values were adjusted across perturbations using the Benjamini–Hochberg procedure, and an adjusted P value below 0.05 was considered significant.

### Neighborhood composition analysis

To determine whether specific non-hepatocyte cell types were spatially enriched or depleted around hepatocytes carrying a given genetic perturbation, we analyzed the cellular composition of local neighborhoods using centered log-ratio (CLR)-transformed compositional data^101^. Within each sample, a local niche was defined for each hepatocyte carrying a single high-confidence perturbation as all neighboring non-hepatocyte cells within a 50-µm radius. Analogous niches were constructed around control hepatocytes.

For each niche, the relative abundance of each neighboring cell type was calculated and transformed using the CLR transformation:

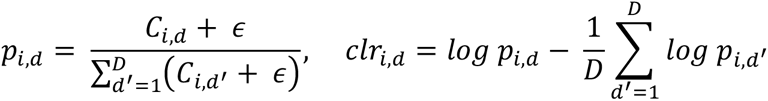

where *i* denotes the index hepatocyte, *d* denotes a neighboring cell type, *C_i_*_,*d*_ is the number of cells of type *d* within the neighborhood of hepatocyte *i*, *D* is the total number of neighboring cell types considered, and *ε* is a pseudocount set to 0.5. *p_i_*_,*d*_ represents the relative abundance of cell type *d* within the local niche of hepatocyte *i*.

For each perturbation and neighboring cell type, CLR values from perturbed niches were compared with those from control niches within each sample using a two-sided Wilcoxon rank-sum test. P values were corrected for multiple testing using the Benjamini–Hochberg false discovery rate (FDR) procedure. A perturbation–cell-type association was considered reproducibly enriched or depleted when a significant change in the same direction relative to control was observed in at least three of five independent datasets, with an adjusted *P* < 0.05 in each qualifying dataset.

For visualization of cross-sample effects, sample-level statistics for each perturbation– cell-type pair were additionally combined across the five datasets using Stouffer’s method to obtain a combined *z* score and corresponding *P* value. Combined P values were subsequently adjusted across all perturbation–cell-type pairs using the Benjamini– Hochberg procedure. These combined FDR values were used only for visualization in the summary dot plots and were not used to define significant perturbation–cell-type associations, which were determined using the reproducibility criterion described above.

### Identification and spatial enrichment analysis of pDC-like cells

To resolve non-hepatocyte subpopulations, each major non-hepatocyte compartment was independently subclustered using its MERFISH expression profiles. Normalized, log-transformed expression values were scaled and clipped at a maximum value of 10, and principal component analysis was performed, retaining 30 principal components. A neighborhood graph was constructed using 15 nearest neighbors, followed by Leiden clustering using the igraph implementation with two iterations and a random seed of 0. The dendritic-cell/monocyte–macrophage compartment was clustered at a resolution of 1.0. Subclusters containing fewer than 300 cells were merged with the nearest subcluster centroid. The resulting 14 subclusters were annotated by marker expression and scoring against curated lineage gene sets. One subcluster containing 4,335 cells showed the highest pDC marker score and was designated pDC-like, reflecting the limited coverage of dendritic-cell identity markers in the targeted MERFISH panel.

Spatial enrichment was quantified using hepatocytes with a single high-confidence sgRNA assignment as anchor cells, with sgRNAs targeting the same gene pooled by target. Hepatocytes carrying non-targeting sgRNAs served as control anchors. Neighboring cells were identified from cell-centroid coordinates using a k-d tree and counted in concentric annuli of 0–15, 15–30, 30–50, 50–75, 75–100, and 100–150 µm. Annular counts were obtained by differencing cumulative counts at successive radii. Each anchor hepatocyte was excluded from its own neighborhood.

For each perturbation, neighboring population, and annulus, enrichment was calculated as the fraction of neighboring cells belonging to that population around perturbed anchors divided by the corresponding fraction around control anchors:

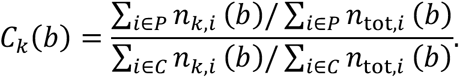

Here, *P* and *C* denote perturbed and non-targeting control anchors, respectively, and *n_k_*_,*i*_(*b*) and *n*_tot,*i*_(*b*) denote the numbers of cells belonging to population *k* and to all populations in annulus *b* around anchor *i*. Counts were summed across anchors and tissue sections before calculating the ratio. This statistic measures relative cellular composition within each annulus, with *C_k_*(*b*) = 1 indicating no enrichment relative to control neighborhoods. Perturbations represented by at least 25 anchors were evaluated and ranked by pDC-like cell enrichment in the 0–15-µm annulus (Fig. 4f). This annular analysis was distinct from the broader distance-dependent neighborhood analysis in Fig. 4g, which summarizes fold changes in neighboring cell fractions across liver samples as mean ± SEM.

Statistical significance was assessed using a label-permutation framework stratified by tissue section. For each population and annulus, we defined *d_i_* = *n_k_*_,*i*_ − *f_C_*_,*s*_*n*_tot,*i*_, where *f_C_*_,*s*_ is the corresponding population fraction around control anchors in section *s*. The test statistic was *S* = ∑*_i_*_∈*P*_ *d_i_*. Under random reassignment of perturbation and control labels within each section, preserving their observed numbers, the mean and variance of *S* were calculated analytically:

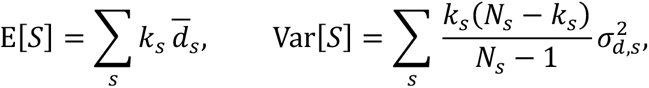

where *k_s_* is the number of perturbed anchors, *N_s_* is the combined number of perturbed and control anchors, and 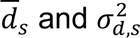 are the mean and population variance of *d_i_* across these anchors in section *s*. Standardized scores were calculated as

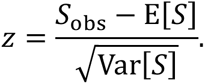

One-sided enrichment P values were obtained using a normal approximation and corrected across all 15,554 perturbation–population pairs tested in the 0–15-µm annulus using the Benjamini–Hochberg procedure.

Enrichment hits were required to satisfy four criteria: an adjusted P value below 0.05; enrichment greater than 1.25 in the 0–15-µm annulus; a negative Spearman correlation between annulus midpoint and enrichment, with maximal enrichment in the innermost annulus; and enrichment greater than 1 in at least four of five tissue sections. Section-specific enrichment was evaluated only when at least 20 perturbed anchors were available in that section.

Confidence intervals for annular distance-dependent enrichment profiles were estimated using 2,000 bootstrap replicates, independently resampling perturbed and control anchors within each tissue section. Neighborhood counts retained anchor multiplicity, such that a cell neighboring multiple anchors contributed to each corresponding neighborhood. Unique-cell counts were calculated separately by assigning each neighboring cell to the annulus containing its nearest anchor.

### Neighborhood differential expression analysis

Spatial neighbors of each perturbed hepatocyte were identified using three-dimensional tissue coordinates in micrometers (x, y, and z) across six cumulative radial windows: 0– 25, 0–50, 0–75, 0–100, 0–125, and 0–150 μm. For each perturbed-hepatocyte instance, neighbor cell type, and distance window, the CP10K-normalized and log1p-transformed expression profiles of all neighboring cells of that type were averaged to generate a single neighbor-expression profile. For each perturbation target and neighboring cell type, profiles associated with gene-targeted hepatocytes were compared with those associated with non-targeting control hepatocytes using a Wilcoxon rank-sum test, separately within each of the five samples. Perturbations represented by fewer than three valid hepatocyte instances were excluded. Within each sample, P values were adjusted using the Benjamini–Hochberg procedure. A gene was classified as a consistent neighbor differentially expressed gene for a given perturbation, neighboring cell type, and distance window if it showed a significant change in the same direction in at least two of the five samples, was not significantly changed in the opposite direction in any sample, and had a cross-sample mean log2 fold change consistent with that direction.

---

**Supplementary Fig. 1.**
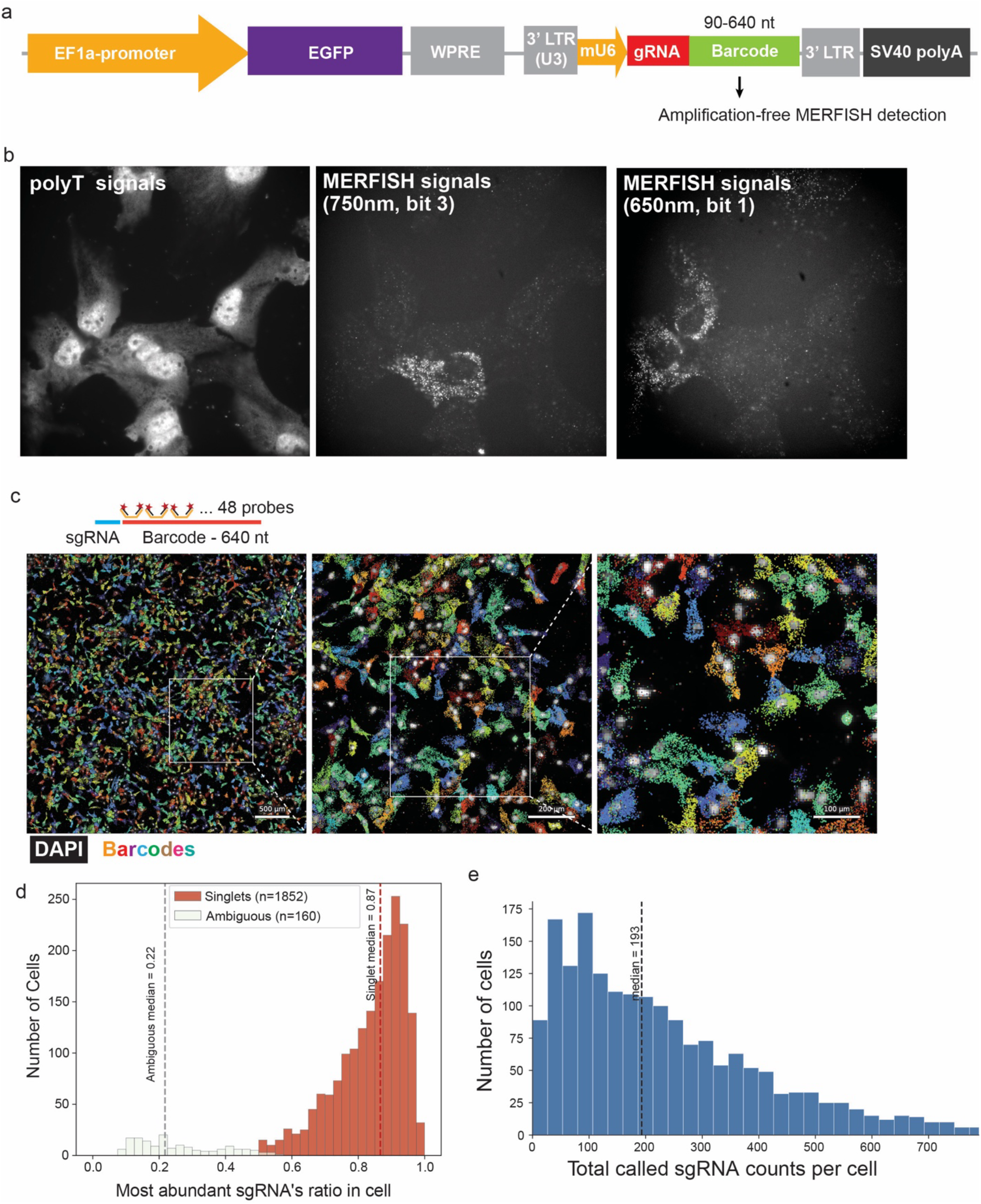
Raw and decoded MERFISH signals from 640-nt sgRNA barcodes. **a,** Schematic of the lentiviral backbone containing the sgRNA–barcode cassette used for *in vitro* Perturb-MERFISH experiments in RPE1 cells. Barcode sequences ranging from 90 to 640 nt were tested in separate experiments for amplification-free detection by MERFISH. For the *in vivo* mouse liver experiments, the promoter and reporter shown here were replaced with a hepatocyte-specific promoter and mTurquoise2, respectively (Materials and Methods). **b,** Representative raw fluorescence images of RPE1 cells infected with lentiviral constructs carrying 640-nt sgRNA barcodes. Poly(T) signal from total mRNA imaging provides visualization of cellular morphology (left). Raw images from two representative MERFISH readout channels (two bits) in the same field of view are shown for bit 3 (750 nm; middle) and bit 1 (650 nm; right). Bright punctate signals are concentrated within subsets of cells, predominantly in the cytoplasm, consistent with cell-specific expression of distinct sgRNA barcodes. **c,** Decoded sgRNA barcodes in RPE1 cells. Distinct sgRNA identities are represented by uniquely encoded barcodes and displayed in different colors overlaid with DAPI-stained nuclei. A global field of view is shown on the left, and two successive magnifications of the indicated regions in the middle and on the right. The schematic above illustrates the 640-nt barcode design and the 48 MERFISH encoding probes targeting each barcode. **d,** Distribution of the fraction of decoded sgRNA molecules assigned to the most abundant sgRNA in single-perturbed cells and cells with ambiguous perturbation assignment. Cells were first filtered to retain those with a segmented volume of at least 800 μm³ and at least three total decoded barcode molecules, including blank barcodes. An sgRNA was called for a cell if it had ≥5 barcode counts and a count fraction ≥0.2; blank barcodes were excluded from calling. Cells were classified as singlets, multiplets, or ambiguous based on these calls. Singlets had exactly one called sgRNA, which accounted for ≥50% of the cell’s total sgRNA counts and had at least three times as many counts as the second-most abundant sgRNA. Multiplets had two or more called sgRNAs, with no additional requirement on their relative abundances. All remaining cells were classified as ambiguous. Histograms show the dominant-sgRNA fraction for singlet and ambiguous cells, with dashed lines indicating the median for each group. **e,** Distribution of the total number of decoded sgRNA-barcode molecules per cell among singlet and multiplet cells defined in **d**. The dashed line indicates the median sgRNA count per cell.

**Supplementary Fig. 2.**
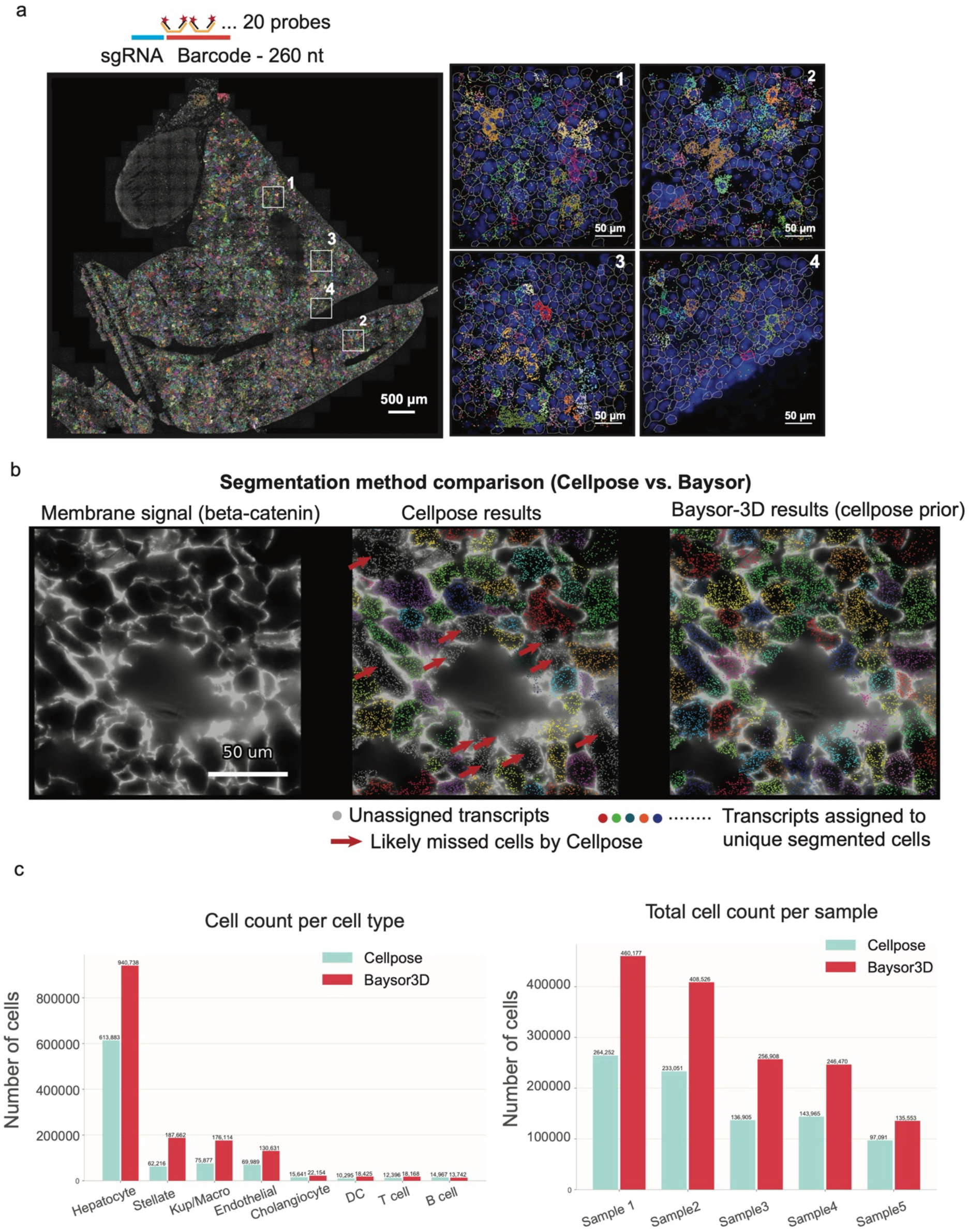
Performance of 260-nt barcode for *in vivo* sgRNA detection and comparison of cell-segmentation methods in mouse liver Perturb-MERFISH data. **a**, Detection of sgRNA barcodes in mouse liver using 260-nt barcodes. Distinct sgRNA identities are represented by uniquely encoded barcodes and displayed in different colors overlaid with DAPI-stained nuclei. A global tissue mosaic is shown on the left, with four magnified views of the indicated regions shown on the right; in the magnified views, DAPI-stained nuclei are shown in blue and cell-segmentation boundaries in white. The upper-left portion of the tissue is likely gallbladder, where few sgRNA barcode signals were detected, consistent with limited activity of the hepatocyte-specific promoter in this tissue. The schematic above illustrates the 260-nt barcode design and the 20 MERFISH encoding probes targeting each barcode. **b**, Comparison of cell segmentation using Cellpose alone and Baysor-3D with Cellpose segmentation masks provided as priors. Membrane-associated β-catenin signal used to guide Cellpose segmentation is shown on the left. For the Cellpose and Baysor-3D results, transcripts assigned to individual segmented cells are shown in distinct colors, whereas unassigned transcripts are shown in gray. Red arrows indicate regions containing abundant transcripts and visually apparent membrane boundaries but no corresponding Cellpose segmentation, consistent with cells missed by Cellpose. Baysor-3D recovered additional cells in these regions by incorporating transcript localization together with the Cellpose prior. **c**, Comparison of cell recovery between Cellpose and Baysor-3D segmentation. Cell counts are shown for individual annotated cell types across the integrated dataset (left) and for each of the five mouse liver samples (right). Baysor-3D recovered more cells than Cellpose alone across most cell types and in all five samples.

**Supplementary Fig. 3.**
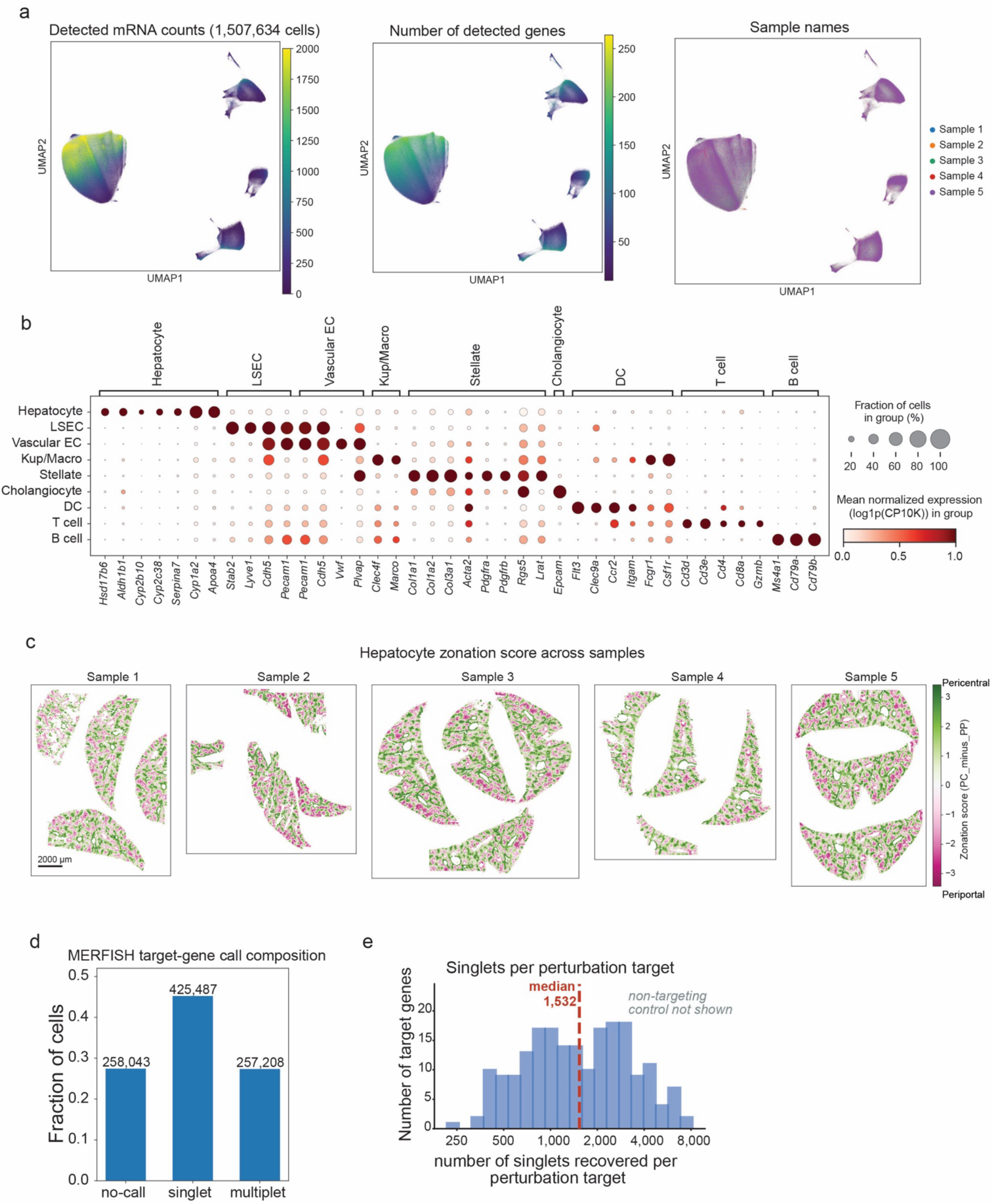
Integrated analysis of Perturb-MERFISH data across five mouse liver samples. **a**, UMAP visualization of the transcription profiles of cells in the integrated Perturb-MERFISH dataset comprising 1,507,634 cells from five mouse liver samples. Cells are colored by the total number of detected mRNA molecules (left), number of detected genes (middle), or sample of origin (right). Data from the five samples were integrated using Harmony (Materials and Methods). **b**, Dot plot showing expression of canonical marker genes across annotated cell types. Dot color indicates mean normalized gene expression within each cell type, calculated as log1p-transformed counts per 10,000 molecules (CP10K), and dot size indicates the fraction of cells with detectable expression of each gene. **c**, Spatial maps of hepatocyte zonation scores across the five mouse liver samples. Positive scores (green) indicate more pericentral-like transcriptional states, whereas negative scores (magenta) indicate more periportal-like states. Zonation scores were calculated from the relative expression of selected pericentral and periportal marker genes (Materials and Methods). **d**, Composition of hepatocytes according to sgRNA assignment. Cells were classified as no-call, singlet, or multiplet based on the number of called perturbations per cell. Bars show the fraction of hepatocytes in each category, with the corresponding cell numbers indicated above the bars. **e**, Distribution of the number of singlet hepatocytes recovered per perturbation target across the five samples combined. Cells assigned to sgRNAs targeting the same gene were pooled by target gene. Among 425,487 singlet hepatocytes assigned to 202 targeted genes, the number of cells recovered per target gene ranged from 234 to 7,587, with a median of 1,532 cells per target. Non-targeting control hepatocytes were excluded from this distribution.

**Supplementary Fig. 4.**
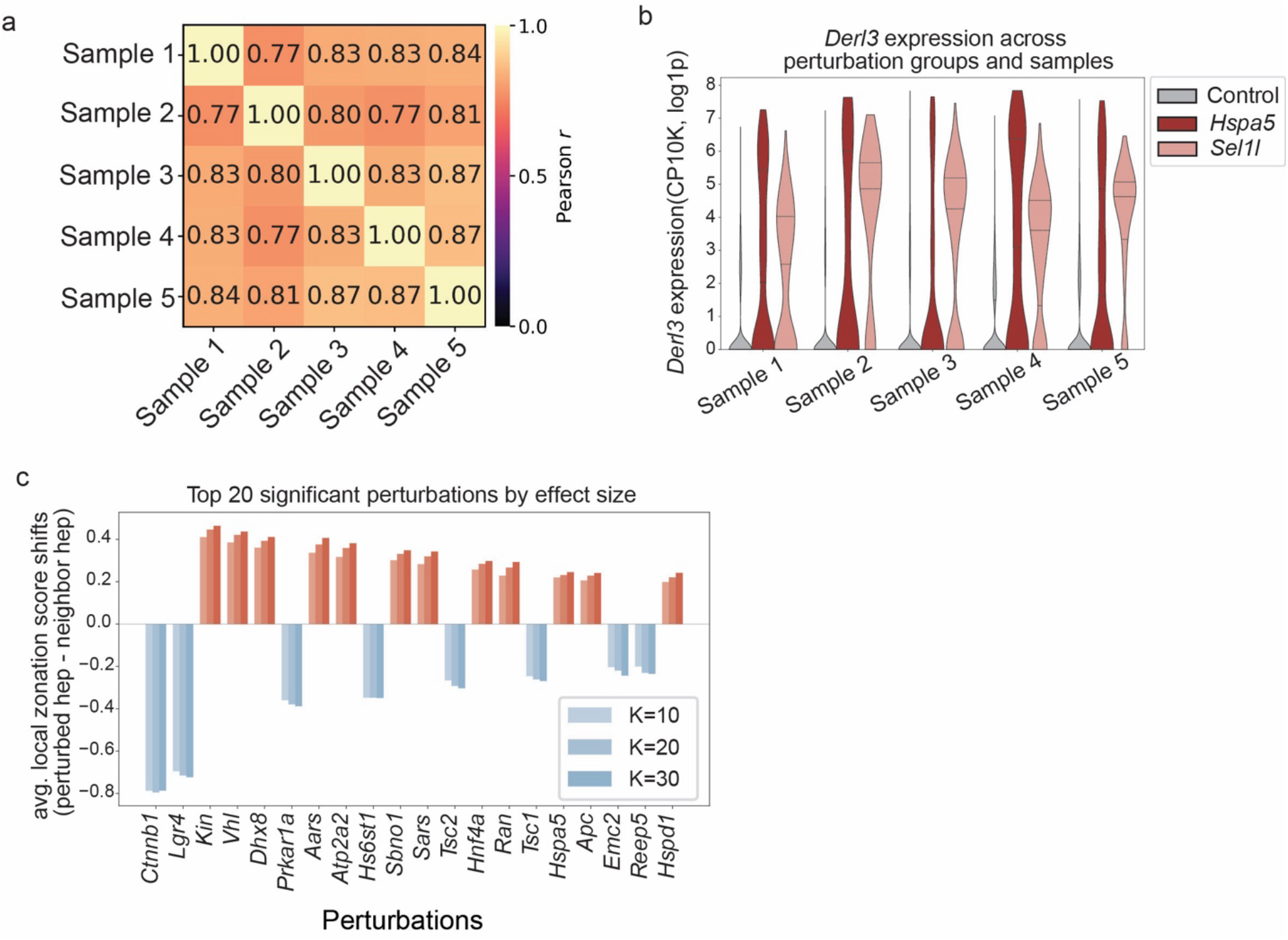
Cell-autonomous perturbation effects across liver samples and zonation shifts of hepatocytes. **a,** Pairwise Pearson correlations between perturbation-effect (transcription profiles) estimates across the five liver samples. **b,** Violin plots showing *Derl3* expression in control, *Hspa5*-targeted, and *Sel1l*-targeted hepatocytes across five independent liver samples. *Derl3* expression was consistently increased following *Hspa5* or *Sel1l* perturbation relative to controls. Expression values are shown as log1p-transformed CP10K; dashed lines indicate the 25th, 50th, and 75th percentiles. **c,** Top 20 significant perturbations producing the largest absolute shifts in hepatocyte zonation scores. Effect sizes were consistent across analyses using different numbers of nearest neighboring hepatocytes (*k* = 10, 20, or 30). Delta was calculated as the difference between the zonation score of each perturbed hepatocyte and the mean score of its *k* nearest neighboring unperturbed hepatocytes (Materials and Methods).

**Supplementary Fig. 5.**
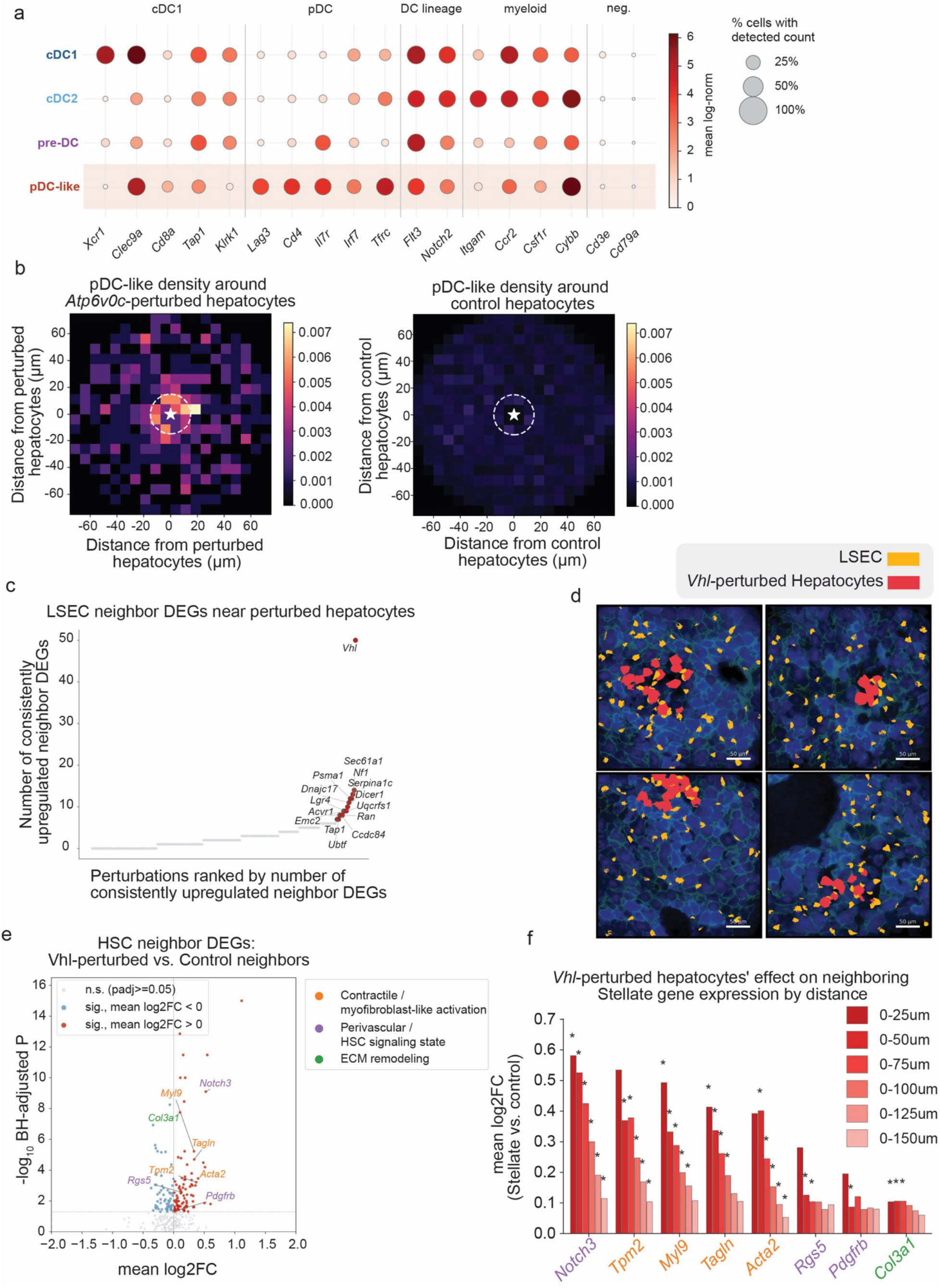
Cell non-autonomous perturbation effects of *Atp6v0c*- and *Vhl*-perturbed hepatocytes. **a,** Expression profiles of representative genes across four dendritic cell subpopulations. Dot color indicates mean expression level, and dot size indicates the fraction of cells expressing each gene. **b**, Two-dimensional density plots of pDC-like cells around *Atp6v0c*-targeted hepatocytes (top; *n* = 1,082) and hepatocytes expressing non-targeting control sgRNAs (bottom; *n* = 19,504). Maps are centered on the anchor hepatocyte, indicated by the white star. **c,** Number of consistently upregulated genes in LSECs within 50 µm of hepatocytes carrying each genetic perturbation. Perturbations are ranked by the number of upregulated DEGs in neighboring LSECs. A gene was considered consistently upregulated if it was significantly increased in at least two of five liver samples, with no sample showing significant regulation in the opposite direction. **d,** Representative images of tissue regions showing local accumulation of LSECs near *Vhl*-targeted hepatocytes. LSECs are shown in yellow and *Vhl*-targeted hepatocytes in red. Scale bars, 50 µm. **e**, Differential gene expression in hepatic stellate cells (HSCs) neighboring *Vhl*-targeted hepatocytes. HSCs within 50 µm of *Vhl*-targeted hepatocytes were compared with HSCs within 50 µm of control hepatocytes expressing non-targeting sgRNAs separately in each liver sample. Log2 fold changes were averaged across samples, and P values were adjusted using the Benjamini–Hochberg method. The x axis shows the mean log2 fold change across samples and the y axis shows −log10 of the minimum adjusted P value across samples. Significant genes are colored red or blue according to the direction of the mean log2 fold change, whereas non-significant genes are shown in gray. Selected genes are annotated according to three manually curated functional programs—contractile/myofibroblast-like activation, perivascular/HSC signaling state, and extracellular matrix remodeling—based on established gene annotations and HSC/pericyte literature. **f**, Distance-dependent expression changes of selected genes in hepatic stellate cells (HSCs) neighboring *Vhl*-targeted hepatocytes. Bars show the mean log2 fold change in HSCs surrounding *Vhl*-targeted hepatocytes relative to HSCs surrounding control hepatocytes expressing non-targeting sgRNAs, within the specified distance, ranging from 25 to 150 µm, of the anchor hepatocyte. Asterisks indicate distances at which the gene was consistently upregulated, defined as significant upregulation in at least two of five liver samples with no sample showing significant regulation in the opposite direction.

